# A molecular framework of chromatin extrusion in plants

**DOI:** 10.64898/2026.01.07.698229

**Authors:** Sofia Tzourtzou, Zhidan Wang, Sarah Leuchtenberg, David Latrasse, Moussa Benhamed, Chang Liu

**Affiliations:** Department of Epigenetics, Institute of Biology, University of Hohenheim, Garbenstrasse 30, 70599 Stuttgart, Germany; Université Paris-Saclay, CNRS, INRAE, Univ Evry, Institute of Plant Sciences Paris-Saclay (IPS2), Orsay 91405, France; Université de Paris Cité, Institute of Plant Sciences Paris-Saclay (IPS2), 91190 Gif-sur-Yvette, France

**Author notes:** Correspondence: Chang Liu.

## Abstract

Three-dimensional genome organization is a fundamental regulator of gene expression, yet the mechanisms shaping higher-order chromatin folding in plants remain poorly defined. Here, we identify the Arabidopsis cohesin regulator PDS5A as a central modulator of chromatin loop extrusion and topologically associating domain (TAD)-like architecture. Using Hi-C and Micro-C, we show that loss of PDS5A strongly enhances TAD-like domains and nucleosome-scale stripes and loops across chromosome arms, while leaving large-scale compartmentalization largely intact and revealing conserved TAD-like patterns across tissues and ploidy levels. Genetic analyses indicate that *Arabidopsis* TAD-like domain formation relies on the plant-specific kleisin SYN4 and is antagonized by WAPL1/2, with PDS5A acting as the dominant cohesin-unloading factor that limits loop extension. Biochemical experiments demonstrate that PDS5A physically associates with cohesin and that its function in suppressing TAD-like domains is independent of direct Tudor domain-mediated binding to H3K4me1-marked chromatin. Micro-C further resolves that chromatin loop and stripe anchors formed in *pds5a* are highly enriched at accessible, strongly transcribed promoters and are marked by the plant site II motif TGGGCC/T, implicating site II-binding transcription factors as plant-specific boundary elements analogous to CTCF in animals. Finally, mutational disruption of distal chromatin anchors shows that newly formed chromatin contacts in *pds5a* can act as cis-regulatory modules that influence target gene expression. Together, these findings identify a PDS5A-regulated chromatin extrusion module that shapes plant 3D genome architecture and uncover a promoter- and motif-based logic for loop anchoring and transcriptional control in plants.

## Introduction

The current understanding of eukaryotic genome organization highlights the profound impact of three-dimensional (3D) chromatin architecture on gene expression, DNA replication, and genome integrity ^1-3^. In this context, topologically associating domains (TADs) represent a structural organizational unit in animal genomes, which are demarcated by increased frequencies of internal chromatin interactions and bounded by regions of strong insulation ^4,5^. Recent advances have revealed that plant genomes, though lacking canonical animal insulator proteins, also exhibit spatial compartmentalization at sub-chromosomal levels, including the presence of so-called "TAD-like" domains ^6-9^. However, mechanisms underlying their establishment and regulation in plants remain poorly understood.

TADs in mammals are contiguous genomic regions featuring high levels of intra-domain contacts and sharp interaction decrease at domain boundaries, visible as triangles along the diagonal in Hi-C contact maps ^5^. TAD boundaries are often enriched for transcription start sites (TSSs), housekeeping genes, active promoters, and specific architectural proteins such as CTCF and cohesin complexes ^4,5,10^. Biochemical and genetic studies in animals have established that some TADs in the genome are essential for regulatory insulation, with their perturbation leading to aberrant gene expression and disease phenotypes ^11-14^.

The formation of animal TADs has been traditionally explained by the loop extrusion model, wherein the ring-shaped cohesin complex loads onto DNA and, powered by ATP hydrolysis, translocates along chromatin to extrude loops of increasing size ^4,15-18^. This process continues until cohesin encounters convergently oriented CTCF binding sites, which serve as a barrier, thereby fixing the boundaries of TADs. The importance of cohesin and CTCF is underscored by genetic depletion experiments: loss of either protein disrupts TAD structure, despite leaving higher-level compartmentalization relatively intact ^19-22^. Additional contributors to domain formation in animals include transcriptional activity and chromatin state, as some boundaries persist even in the absence of CTCF or cohesin ^23-26^. In yeast, which lacks CTCF, cohesin-mediated loop extrusion can be terminated by collision with other cohesin complexes rather than sequence-specific barriers, highlighting potential alternative mechanisms conserved among eukaryotes. In this scenario, cohesin-associated regions, which are often located at TTSs of convergently transcribed genes, act as stalling points for extrusion ^27^.

Plant genomes display complex and diverse 3D chromatin organizations. TAD-like domains have been characterized in crops with large genomes (e.g., rice, sorghum, maize, tomato, and cotton), as well as in small-genome species like *Marchantia polymorpha* ^7,28,29^. Plant TAD-like domain boundaries are often enriched in transcriptionally active regions and active histone marks; however, typical long-range border-to-border contacts, which are hallmarks of animal TADs, are less pronounced. In the dicot model plant *Arabidopsis thaliana,* which has a smaller genome compared to other plants, TAD-like domains are weak and largely unrecognizable. In the *Arabidopsis* genome, only a limited number of domain-like structures were detected, aligning with epigenetic compartmentalization ^30-32^. These observations have fostered debate about whether plant TAD-like domains arise from the same regulatory mechanisms as animal TADs ^33-35^.

In plants, clear CTCF homologs are absent, yet core cohesin components and candidate regulators are conserved ^24^. Several plant-specific DNA- and protein-based barriers have been proposed to modulate domain boundaries, such as regulators of Polycomb group proteins (such as PWO1 and EMF1) and TCP transcription factors ^29,31,32,36,37^. However, disruption of these proteins has not led to the complete loss of TAD-like demarcation, suggesting further mechanistic complexity. Superimposed on these potential boundary factors, changes in epigenomic modifications can modulate local domain formation in plants, but are often insufficient to fully explain TAD-like architecture ^38-41^.

Recently, we found that mutation of PDS5A, which encodes a coregulator of cohesin, remarkably enhanced TAD-like structures in the *Arabidopsis* genome ^42^. This finding raised fundamental questions about the evolution of chromosome architecture in eukaryotes and the extent to which transcriptional activity, epigenomic states, and mechanical constraints jointly shape domain organization in plant nuclei. Here, we present a systematic identification of cohesin subunits and regulatory factors that govern TAD-like domain formation in *Arabidopsis thaliana*. By combining targeted genetic perturbations with advanced chromosome conformation capture-based approaches, we delineate how cohesin and its cofactors shape higher-order chromatin architecture and assess the degree to which these mechanisms converge with or diverge from paradigms established in animals. Furthermore, the identification and functional characterization of plant-specific domain boundaries and their regulators provides new insights into the general principles underlying plant genome folding.

## Results

### PDS5A-dependent TAD-like domain architecture in Arabidopsis is conserved across tissues and ploidy levels

In our recent study, we demonstrated that PDS5A functions as a key regulator of chromatin organization in *Arabidopsis* via suppressing the formation of TAD-like domains at the genomic level. To further elucidate the mechanisms underlying TAD boundary establishment in the *pds5a* mutant, we investigated several potential contributing factors, such as transcriptional reprogramming, endopolyploidy levels, and genetic interactions.

We began by comparing Hi-C maps generated from leaf, root, and germinating seed tissues, which represent distinct developmental stages and gene expression profiles. As expected, numerous genomic regions were clearly identifiable as TAD-like domains in *pds5a* leaf tissues, whereas these domains were hardly recognizable in the corresponding Hi-C maps of wild-type samples (Fig. 1a, supplemental Tables 1 and 2). The enhancement of TAD-like domains in *pds5a* is accompanied by the strengthening of local chromatin insulation ^42^. Insulation scores were calculated to quantify potential changes in chromatin insulation across samples ^43^, which indicated that chromatin insulation in leaves and roots was highly similar, whereas it was weaker in germinating seeds (Fig. 1b,c). Aggregate TAD plots, which depict normalized observed/expected chromatin contacts within and across TAD-like domains, revealed that these structures remained unchanged in roots but were weakened in germinating seeds (Fig. 1d). Besides, we examined the patterns of TAD-like domains in leaves of heat-stressed plants, as heat stress is known to induce changes in chromatin organization at a chromosomal level, for instance, dissociation of KNOT ENGAGED ELEMENT (KEE) regions that form megabase-sized chromatin loops ^44,45^. As expected, in heat-stressed *pds5a* leaves, we observed complete dissociation of KEEs (Supplemental Fig. 1). Remarkably, despite such chromatin reorganization, TAD-like domains at the local level were well maintained, suggesting that interactions mediated by KEEs in *Arabidopsis* are independent of *PDS5A* (Supplemental Fig. 1b). Given that the above-mentioned tissues display substantial transcriptomic differences compared to untreated leaf samples (Supplemental Table 3), we further investigated whether differentially expressed genes (DEGs) in various tissues preferentially localize within, outside, or at the borders of TAD-like domains. Our analysis revealed no significant preference for DEGs with respect to their location at TAD-like domains (Fig. 1e). Likewise, chromatin regions containing DEGs did not show significant alteration in their insulation scores (Supplemental Fig. 2). These results are in agreement with previous studies that induced alterations in animal TADs genome-wide, yet showed only marginal changes in gene expression ^21,46-48^.

**Figure 1.**
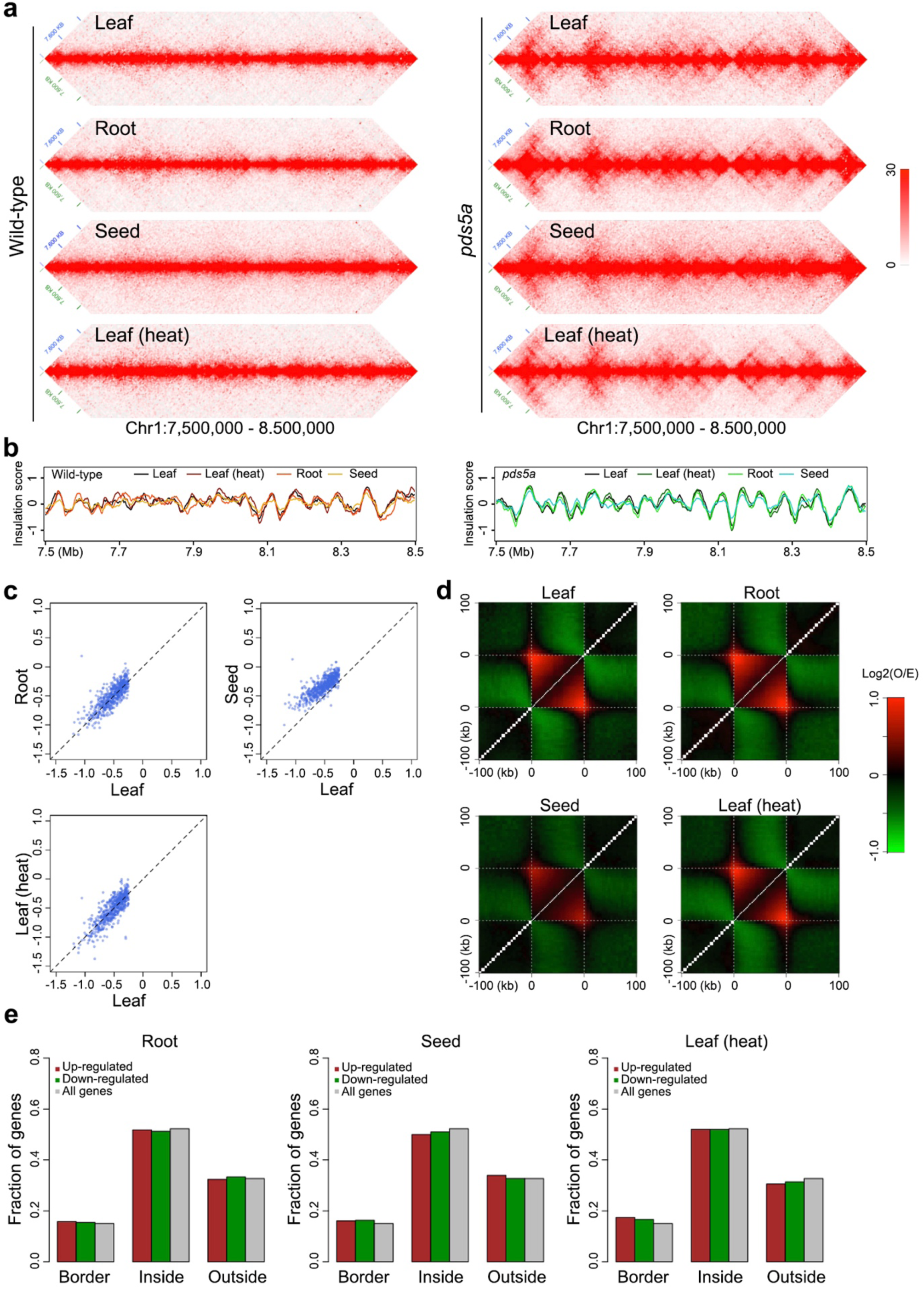
Comparative analysis of chromatin insulation, TAD-like domains, and gene expression in *pds5a*. **a, b,** Comparison wild-type and *pds5a* Hi-C contact maps (a) and the corresponding profiles (b), both calculated at 5 kb resolution for a 1 Mb region of chromosome 1. Regions with local minima in the insulation score indicate positions of strong chromatin insulation. **c,** Correlation of genome-wide insulation scores between different pds5a samples. Scatter plots display insulation scores specifically from regions exhibiting strong chromatin insulation. **d,** Metagene plot depicting relative chromatin contact strengths within *pds5a* TAD-like domains and their 100 kb flanking regions. Relative chromatin contact was calculated as the log2 ratio of observed versus expected contact frequencies (log2(O/E)), based on normalized Hi-C matrices. **e,** Genomic distribution of differentially expressed genes in relation to TAD-like domains in pds5a. "Border," "Inside," and "Outside" indicate genes located at TAD-like domain borders, within TAD-like domains, and in the regions between TAD-like domains, respectively. "All genes" serves as a background reference and includes all genes in the genome.

Endoreduplication, a process in which cells replicate their genomic DNA without undergoing subsequent cell division, is a common phenomenon in *Arabidopsis* ^49,50^. In 2-week-old *Arabidopsis* seedlings that grow under standard conditions, approximately half of the nuclei had undergone at least one endocycle (Supplemental Fig. 3a). To clarify whether *psd5a* “TAD-like domain” patterns were attributed to sister chromatid contacts in endoreduplicated nuclei, we generated endopolyploidy-specific Hi-C maps ^51^. Our results demonstrated that 2C nuclei (i.e., without endoreduplication) had similar TAD-like domain patterns compared to 4C and 8C nuclei, excluding the participation of endoreduplication-specific mechanisms underlying TAD-like domains formation in *pds5a* (Supplemental Fig. 3b). Taken together, our results indicate that TAD-like domains in *Arabidopsis* are conserved throughout the plant body and are largely independent of changes in gene expression.

### WAPL1/2 and SYN4 participate in regulating TAD-like domains

Constituent members of cohesin and its co-regulators that modulate cohesin-chromatin interaction dynamics are highly conserved in eukaryotes ^52^. Via extruding chromatin, the cohesin protein complex plays a pivotal role in defining TAD structures in animals ^53^. In plants, WAPL acts as a cohesin release factor that promotes cohesin dissociation from chromatin, thereby regulating cohesin dynamics and preventing excessive chromatin loop extension; while SYN2 and SYN4 act as cohesin subunits in somatic cells, known for maintaining proper cohesin function ^54,55^. To examine genetic interactions with *pds5a*, we generated higher-order mutant combinations and analyzed their chromatin architecture by Hi-C.

Compared with *pds5a* alone, *pds5a wapl1 wapl2* and *pds5a syn4* mutants displayed strikingly distinct intrachromosomal contact patterns. Loss of WAPL activity in the *pds5a* background caused a strong reinforcement of long-range chromatin interactions that extended across TAD-like domain boundaries, consistent with unrestrained loop extrusion (Fig. 2a). In contrast, *pds5a syn4* mutants completely lacked detectable TAD-like domains, indicating a failure to establish higher-order chromatin organization (Fig. 2a). Notably, depletion of WAPL proteins in animal cells produces comparable alterations in chromatin contact profiles ^19,20^, underscoring a conserved role for WAPL in constraining cohesin activity (Supplemental Fig. 4). On the other hand, *syn2* mutation did not alter TAD-like domains in *pds5a* (Fig. 2a). These divergent effects on *pds5a* were also reflected by chromatin contact strength curves plotted against genomic distance. Specifically, *WAPL* mutations markedly enhanced long-range chromatin interactions in *pds5a*, whereas introduction of *syn4* restored the chromatin contact profile of *pds5a* to wild-type level (Fig. 2b). Altogether, our results indicate that cohesin-mediated chromatin extrusion is dependent on *SYN4* (a plant-specific kleisin homolog) but is suppressed by *PDS5A* and *WAPLs*.

**Figure 2.**
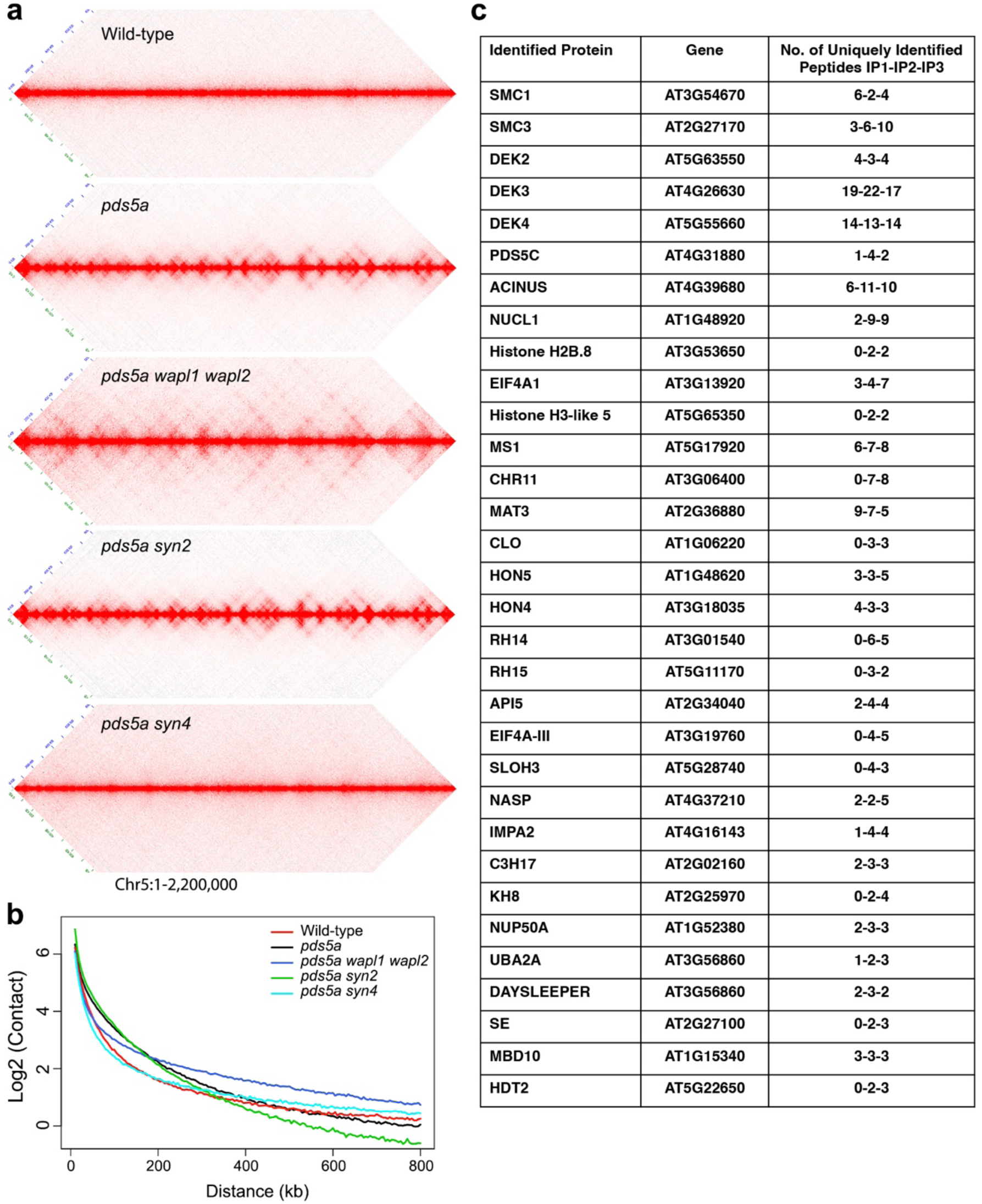
Involvement of cohesin in *Arabidopsis* chromatin loop extrusion. **a,** Comparison of Hi-C maps of different genotypes. The graphs illustrate Hi-C maps of a representative genomic region (2.2 Mb from chromosome 5), normalized at 5 kb resolution. **b,** Decay of chromatin contact frequency along with increasing genomic distance. **c,** Protein interaction partners of PDS5A. PDS5A:2HA proteins were pulled down, and the associated interacting proteins were identified by mass spectrometry. Wild-type (WT) plants were used as controls. The table shows proteins identified in at least two replicates but not in WT.

In non-plant species, PDS5 interacts with cohesin to promote cohesin release from chromatin as well as to prevent cohesin from establishing TAD boundaries ^56,57^. Consistent with this role, proteomic analyses demonstrate that *Arabidopsis* PDS5A directly associates with cohesin: immunoprecipitation of PDS5A:2HA robustly recovered the core cohesin subunits SMC1 and SMC3 (Fig. 2c), and SYN4 was also detected (Supplementary Table 4). Given the essential role of SYN4-containing cohesin complexes in TAD-like domain formation, these findings identify PDS5A as a central negative regulator of extrusion-competent cohesin in plants.

#### Functional dissection of PDS5A domains

The *Arabidopsis* PDS5 family comprises five homologs, yet PDS5A stands out due to its specific role in regulating chromatin organization ^42,58^. In particular, PDS5A and PDS5B are close homologs with highly similar domain architectures. Both proteins harbor arrays of Armadillo domains, consisting of multiple fold structures that together adopt a superhelical conformation; in addition, both contain a Tudor domain near their C-termini ^59,60^. To dissect the molecular basis of PDS5A-specific function, we generated several chimeric fusion constructs in which domains of PDS5A were partially replaced with the corresponding regions from PDS5B (Fig. 3a). All constructs were expressed under the control of the native PDS5A promoter and introduced into the *pds5a* mutant background. Hi-C analysis of these transgenic plants revealed that the N-terminal Armadillo domain array of PDS5A played a minor role in suppressing TAD-like domains, as a fusion protein lacking this region was partially capable of suppressing TAD-like structures (Fig. 3b,c). In contrast, the central Armadillo domain array and the C-terminal region containing the Tudor domain of PDS5A were essential, as replacing these segments with those from PDS5B yielded chimeric proteins that failed to rescue the *pds5a* phenotype, with TAD-like domains remaining unchanged (Fig. 3b,c). Together, these domain-swap experiments suggest that the Armadillo array and the C-terminal Tudor-containing region of PDS5A jointly confer its unique capacity to control higher-order chromatin organization in *Arabidopsis*.

**Figure 3.**
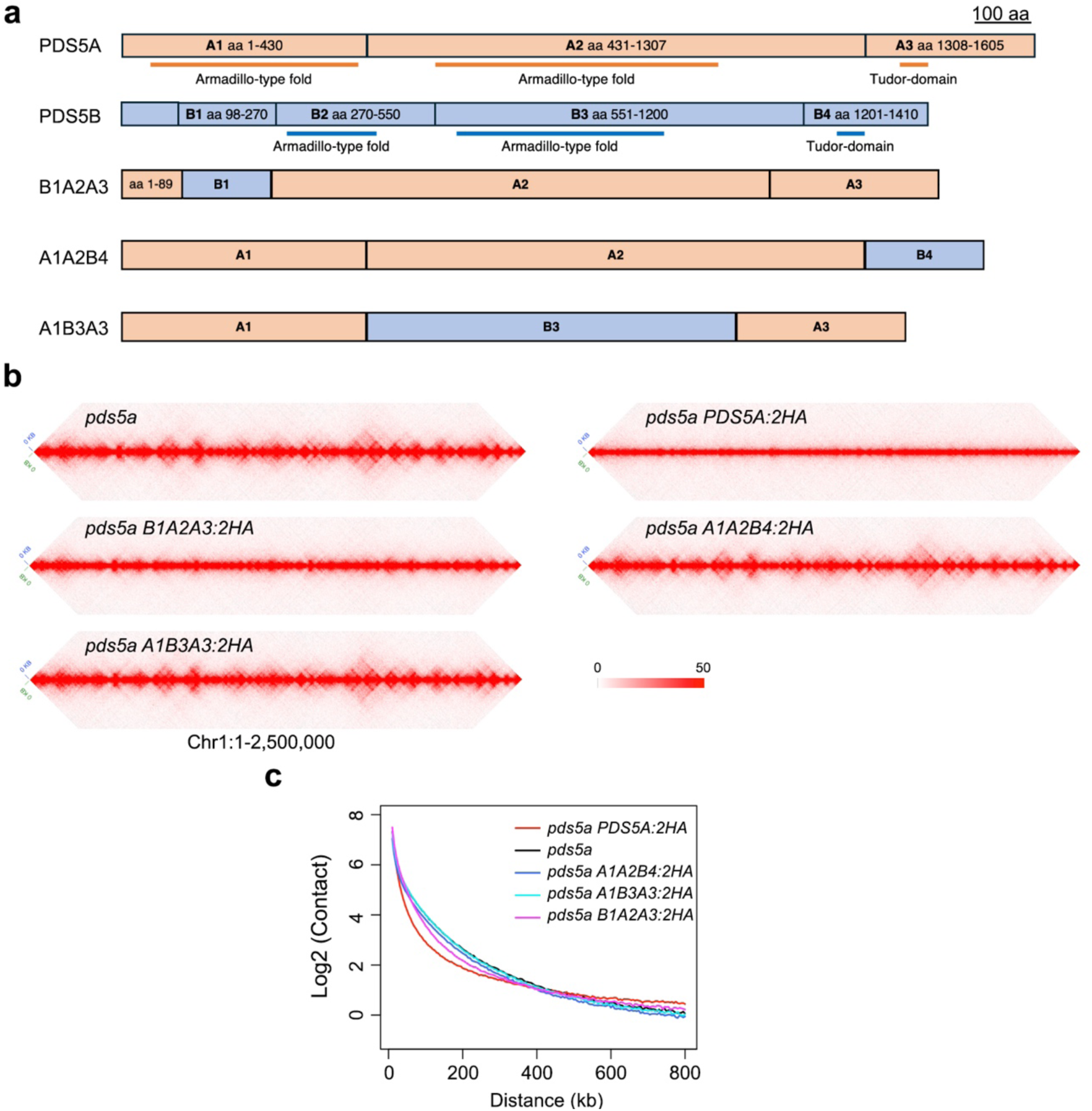
All domains in PDS5A are jointly required to regulate TAD-like domains. **a,** Illustration showing the fragments of PDS5A and PDS5B used for the creation of chimeric fusion proteins exchanging the armadillo-type folds and Tudor-domain of PDS5A with fragments of PDS5B. Domain features highlighted below PDS5A and PDS5B are based on sequence analysis at InterPro (https://www.ebi.ac.uk/interpro/). **b,** Comparison of Hi-C maps of different genotypes. The graphs illustrate Hi-C maps of a representative genomic region (2.5 Mb from chromosome 1), normalized at 5 kb resolution. **c,** Decay of chromatin contact frequency along with increasing genomic distance.

PDS5A was proposed to regulate genome organization via binding to chromatin with its Tudor domain, which recognizes methylated histone H3K4 (H3K4me1) ^42^. Indeed, our ChIP-seq experiment using PDS5A:2HA revealed interactions between PDS5A and H4K4me1-marked chromatin regions (Fig. 4a). Interestingly, our inspection on PDS5A-chromatin interactions around TAD-like borders, which show enhanced chromatin insulation in *pds5a* mutants, did not reveal specific patterns (Supplemental Fig. 5). To verify whether the Tudor domain is required by PDS5A to interact with chromatin, we created a construct containing a *PDS5A:2HA* variant (*PDS5A:2HA^3A^*), in which three amino acid residues in the Tudor domain and likely crucial for interactions with H3K4me1 were mutated ^61,62^. This construct was introduced to the *pds5a/b* genetic background, in which the *PDS5A:2HA^3A^* variant could achieve protein abundance comparable to that in the complementation line (Fig. 4b). Clearly, ChIP-seq experiments indicated that mutated PDS5A:2HA could not bind to H3K4me1-marked chromatin (Fig. 4a), confirming that the Tudor domain is essential for PDS5A’s chromatin association *in vivo*. We next employed Hi-C analysis to determine whether the mutated PDS5A:2HA retains its ability to regulate chromatin organization. To our surprise, PDS5A:2HA^3A^ protein fully suppressed the TAD-like domain structures in *pds5a/b* to wild-type patterns, indicating that its function in chromatin organization remains intact (Fig. 4c,d). Together, our results demonstrate that PDS5A suppresses TAD-like domain formation independently of direct chromatin binding.

**Figure 4.**
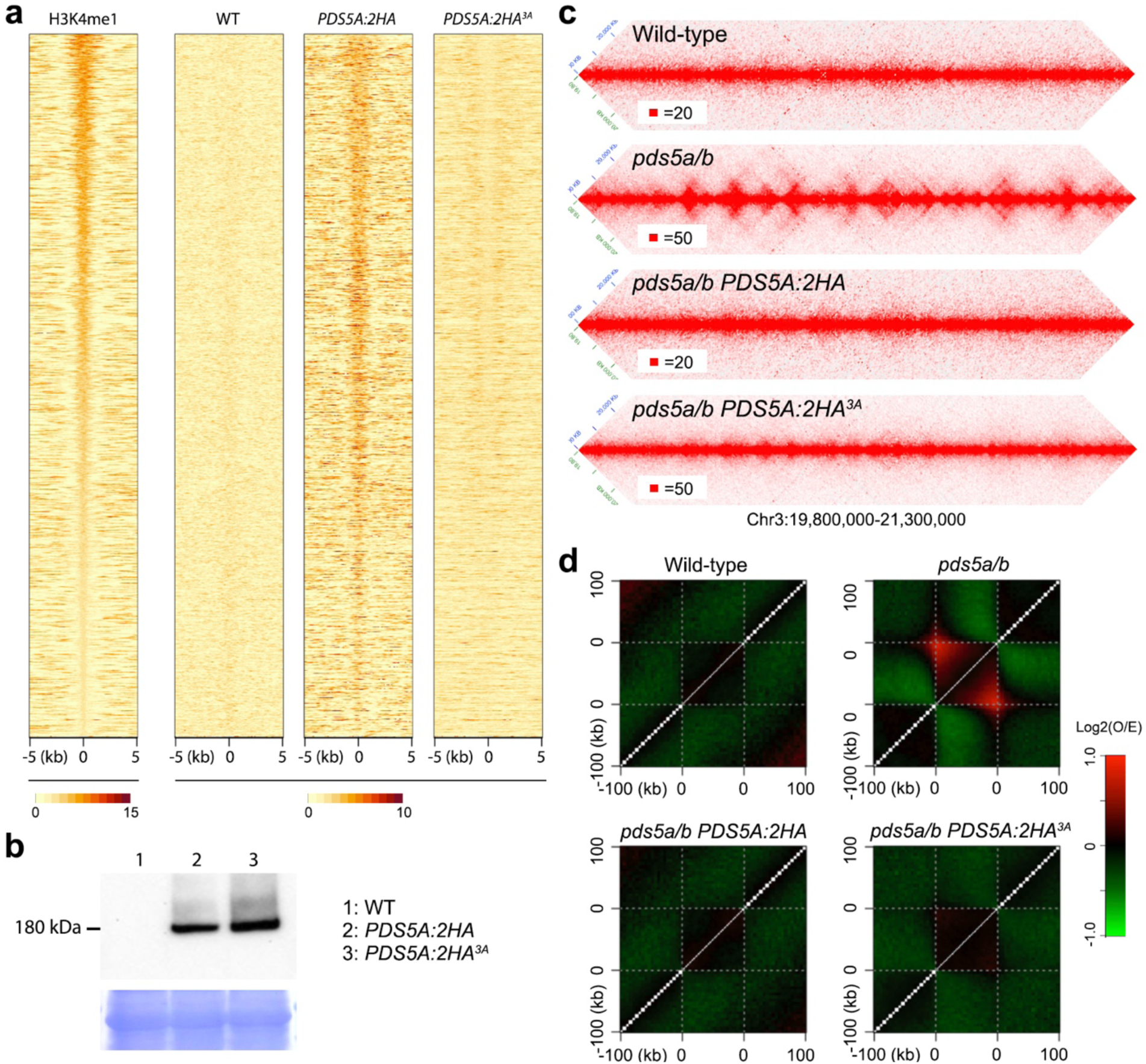
PDS5A regulates TAD-like domain formation independently of interacting with chromatin. **a,** Normalized ChIP-seq reads distribution around chromatin regions marked with H3K4me1. **b,** Protein expression of *PDS5A:2HA* and *PDS5A:2HA^3A^*. Total protein from shoots of 14-day-old WT, *PDS5A:2HA* and *PDS5A:HA^3A^* plants was analyzed by western blot, the membrane was probed with anti-HA antibody and stained with Coomassie Blue to verify equal loading of total protein across lanes. A band corresponding to PDS5A was detected in both tagging lines at the expected size of 180 kDa. **c,** Comparison of Hi-C maps of different genotypes. The graphs illustrate Hi-C maps of a 1.5 Mb representative genomic region, normalized at 5 kb resolution. **d,** Metagene plot depicting relative chromatin contact strengths within TAD-like domains and their 100 kb flanking regions. Relative chromatin contact was calculated as the log2 ratio of observed versus expected contact frequencies (log2(O/E)), based on normalized Hi-C matrices.

#### Fine-scale chromatin stripes and loops emerge at promoters

On mammal Hi-C maps, two prominent signatures of loop extrusion are “corner peaks” (enriched focal contacts at TAD borders or loop anchors) and linear “stripes” extending inward from these peaks, reflecting cohesin-driven loop formation and CTCF-defined barriers ^4^. To investigate fine-scale chromatin contact patterns in *pds5a*, we employed the Micro-C method ^63,64^, which enables the capture of genome-wide DNA interactions at a nucleosomal resolution. When analyzed at a resolution of 1 kb or higher, we found that the *pds5a* Micro-C map displayed widely distributed stripes and dots, which were barely observable in wild-type Micro-C data (Fig. 5a). These features, which largely overlapped (Fig. 5b), were predominantly localized at TAD-like domain boundaries, suggesting that they emerge from the dynamic interplay of cohesin-mediated loop extrusion and the presence of boundary elements that constrain loop expansion.

**Figure 5.**
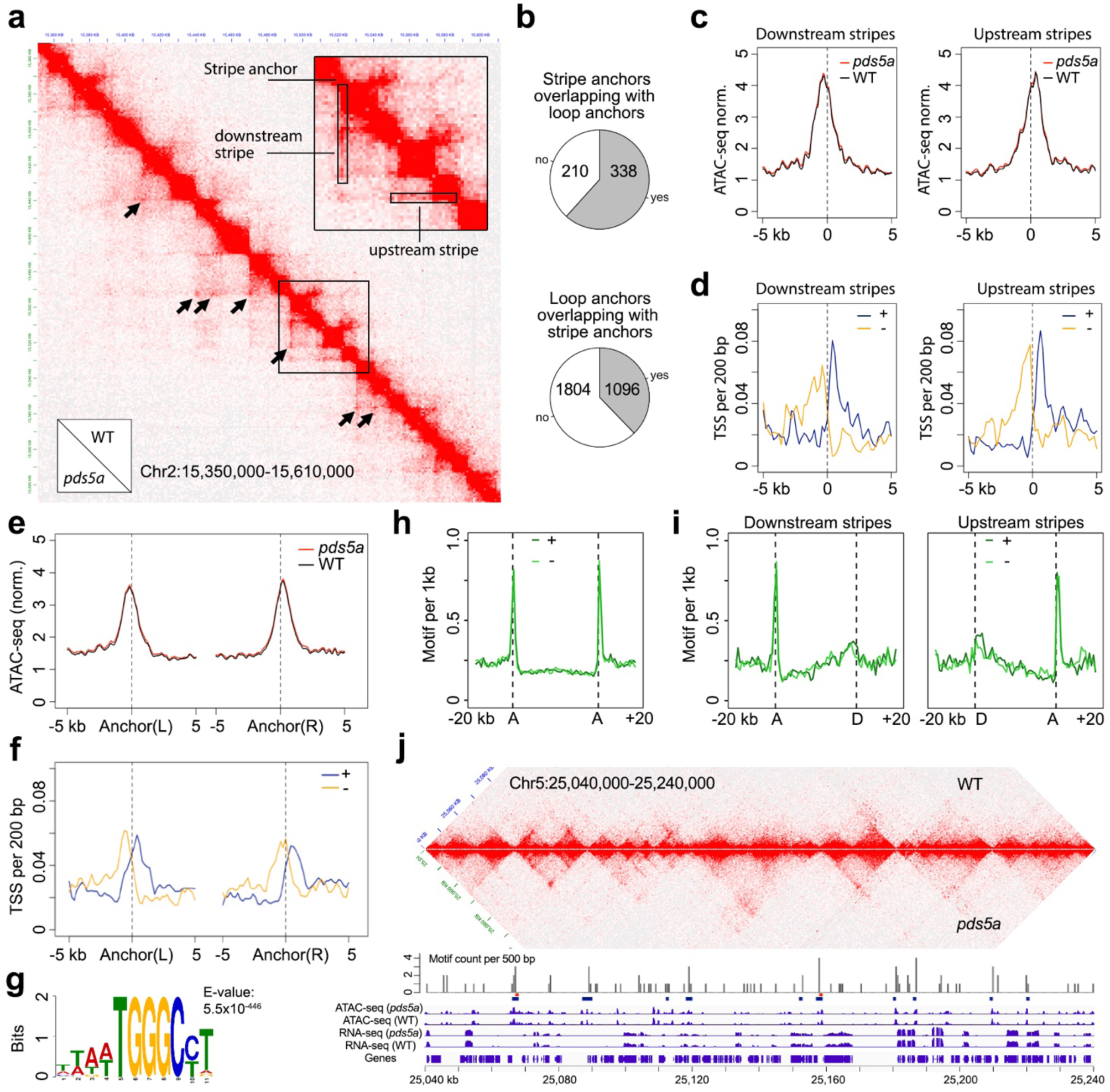
Characterization of stripes anchor and chromatin loop anchor regions. **a,** Example showing stripes (inset) and chromatin loops (indicated by arrows) as detected in the *pds5a* Micro-C map normalized to 1 kb resolution. **b,** Substantial overlap between stripe anchor regions and loop anchor regions. **c,** Comparison of chromatin accessibility at stripe anchors in *pds5a* mutant and wild-type samples, shown with normalized ATAC-seq read coverage. d, Density of transcription start sites (TSS) per 200 bp at stripe anchors, shown separately for plus-strand (blue) and minus-strand (orange) transcripts. **e,** Chromatin accessibility at loop anchors in *pds5a* mutant and wild-type samples, presented as normalized ATAC-seq read coverage. For each chromatin loop, the interacting regions are labeled as L (left) and R (right) according to their genomic coordinates. **f,** Density of TSS per 200 bp at loop anchors, separated by plus-strand (blue) and minus-strand (orange) transcripts. **g,** Sequence logo representing the overrepresented motif identified at loop anchor regions. **h, i,** Density profiles showing motif enrichment (motifs per 1 kb) around loop anchors (h) and stripe regions (i). The presence of the site-II motif (5’-TGGGCC/T-3’) is shown for the plus strand (dark green) and minus strand (light green) separately. In these plots, anchor regions and stripe region termini are labeled as "A" (anchor) and "D" (distal end), respectively. **j,** Integrated overview of stripes and loops within a representative 200 kb genomic region, displayed alongside tracks for various genomic features. The motif count track indicates the number of site-II motifs on both strands. Blue and red segments below the motif count track denote loop anchor and stripe anchor regions, respectively.

To systematically identify and analyze stripes, we implemented an adapted version of the Zebra algorithm, which was originally developed for mammalian Hi-C data ^18^ (see Methods for details; Supplemental Fig. 6a). Applied to 1-kb resolution Micro-C maps of pds5a samples, this pipeline identified 548 stripes ranging from 1 to 4 kb in length (Supplemental Fig. 6b). Each stripe originated from an “anchor” region and typically extended 30-40 kb into the adjacent “stripe domain” (Supplemental Fig. 6c). Integration with *Arabidopsis* epigenomic datasets revealed that stripe anchors co-localized with regions of high chromatin accessibility (Fig. 5c). Furthermore, these anchors were primarily situated near outward-facing transcription start sites (TSS) of genes that were preferentially expressed at high levels (Fig. 5d; Supplemental Fig. 6d).

In the *pds5a* Micro-C maps, distinct dots represented chromatin loops, corresponding to pairwise chromatin contacts between specific genomic loci, which we termed loop anchors. Chromatin loops were identified from the same 1-kb resolution Micro-C maps (see Methods; Supplemental Table 2). Notably, loop anchors shared several features with stripe anchors, including localization within highly accessible chromatin regions and enrichment at promoters of genes with high expression levels (Fig. 5e,f; Supplemental Fig. 6d).

In mammals, chromatin extrusion is halted by transcription factor CTCF at specific loci that contain CTCF motifs ^17,65,66^. From *pds5a* micro-C data, a particularly interesting finding was the strong enrichment of the TGGGC(C/T) motif, known as the plant "site II" element, at stripe anchors and loop anchors (Fig. 5g-j). This was also the only motif revealed by our de novo sequence analysis. The site-II motif is enriched in *Arabidopsis* promoter regions, and it has in vivo interactions with members of several types of transcription factor families including TCP, TRB, and LBD ^67-69^. The striking co-localization of loop and stripe anchors with the plant-specific site II motif implies that this sequence element, along with its associated transcription factors, functions as a candidate regulatory factor that may act analogously to mammalian CTCF by demarcating boundaries or stabilizing loop anchors during cohesin-mediated loop extrusion.

#### Distal promoters function as enhancers under extended chromatin looping

Because *pds5a* broadly strengthens intra-chromosomal contacts among highly accessible promoters, we asked whether distal promoters in cis might act as novel cis-regulatory elements. We therefore selected three *pds5a*-upregulated genes whose promoters formed new cis contacts with distal loci, which we designated as potential enhancers (Fig6. a-c, Supplemental Figure 7). Mutant lines harboring T-DNA insertions adjacent to these candidate enhancers and within the loop-forming region were crossed with *pds5a* to test their impact on target gene expression.

For all these three genes, *pds5a*-dependent up-regulation was suppressed by the T-DNA insertion (Fig. 6d). However, for *AT2G38820*, the T-DNA insertion next to the potential enhancer caused strong up-regulation, potentially masking any additional *pds5a-*dependent effect, so this gene was not analyzed further. For the other two genes, we performed Capture Micro-C to map their chromatin interaction networks in different genetic backgrounds (Supplemental Figure 8). As expected, since T-DNA increased the physical distance between loci that span the insertion site, a strong decrease in chromatin contacts over the T-DNA locus was observed; other than that, we did not find drastic changes in chromatin contact patterns in Capture Micro-C probed regions (Fig. 6e,g). In particular, T-DNA insertion adjacent to their potential enhancers did not disrupt the respective chromatin loops (Fig. 6e,g). Next, we examined promoter chromatin contact in detail by extracting their Micro-C contacts. Interestingly, independent of *PDS5A*, the T-DNA insertion itself did not alter promoter–enhancer contact patterns. (Fig. 6f,h). It is worth noting that, in these two cases, the T-DNA insertions are located approximately 15 kb and 25 kb away from the respective target genes, which is much farther than the expected median distance of ∼1 kb between enhancers and their target promoters in *Arabidopsis* ^70,71^. These T-DNA insertions changed gene expression only under chromatin loop formation (Fig. 6d), demonstrating that enhanced chromatin extrusion in *pds5a* can extend the functional range of cis-regulatory communication much beyond canonical promoter-enhancer distances.

**Figure 6.**
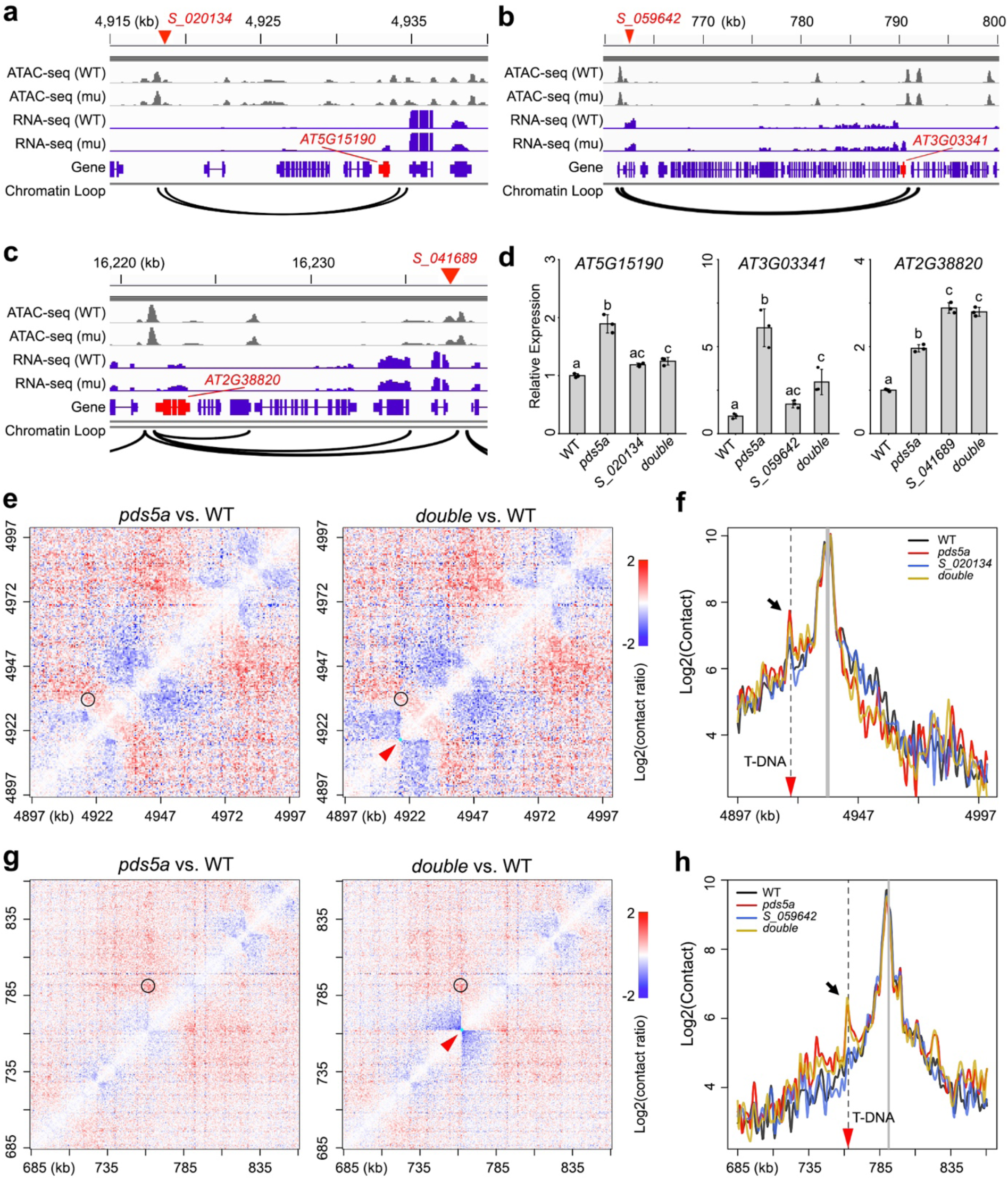
Newly established cis-chromatin contacts in *pds5a* influence gene expression. **a-c,** Genomic regions containing *pds5a*-dependent chromatin loops (arcs) whose anchor regions overlap with genes that are upregulated in *pds5a* (gene models in red). Gene expression and chromatin accessibility in these regions are shown by RNA-seq and ATAC-seq tracks, respectively. WT, wild type; mu, *pds5a*. In each panel, the red triangle marks the insertion site of the SALK line used in this study. **d,** Gene expression in various genetic backgrounds, expressed as mean +/- standard deviation of three biological replicates. For each set of comparisons, “double” refers to the double mutant. Different letters above the bars indicate statistically significant difference at p<0.05 determined by one-way analysis of variance followed by Tukey’s post hoc test. **e,** Comparison of Micro-C contact maps at 500 bp resolution. All maps were generated from Capture Micro-C data, and the full genomic region covered by the capture probes is shown. The red arrowhead points to the T-DNA insertion site; the circle highlights the chromatin loops indicated in panel a. **f,** Chromatin contacts between the promoter (marked by a gray vertical bar) and its surrounding regions, derived from Capture Micro-C. The arrow indicates the chromatin loops described in panel a. The dotted line, together with the red arrowhead, marks the T-DNA insertion site. Genotype labels are as in panel d. **g,h,** As in panels e and f, respectively, but for the genomic region containing the chromatin loops described in panel b.

## Discussion

In this work, genetic and biochemical data establish *PDS5A* as a key negative regulator of cohesin-dependent chromatin extrusion in somatic *Arabidopsis* cells. Loss of *PDS5A* markedly strengthens local insulation and TAD-like domains and induces abundant short-range stripes and focal loops, which are further enhanced by *WAPL1/2* loss and are fully dependent on *SYN4* (Fig. 2). This chromatin extrusion regulatory system resembles that in animals, where *WAPL* depletion substantially extends chromatin loop length ^19^. However, in *Arabidopsis*, *PDS5A* clearly dominates over *WAPL*: mutations in *WAPL* alone have little impact on TAD-like domains, whereas pds5a already suffices to reveal a robust loop-extruded layer, and *pds5a wapl1 wapl2* further extends long-range arm contacts (Fig. 2b). Together with the physical association of PDS5A with cohesin SMC1/3 (Fig. 2c), these findings support a model in which PDS5A is the major unloading factor that sets an upper bound on loop extension in plant chromosome arms.

The strong genetic requirement for *SYN4* places this plant-specific kleisin at the core of the extrusion module uncovered in *pds5a*. *SYN4* is absent outside plants and contains distinctive low-complexity segments compared to canonical mitotic kleisins^72^. Recent work on *syn4* mutants shows broad alterations of intra- and inter-chromosomal contacts, particularly at telomere-proximal regions, and impacts on dynamic stress-responsive gene networks ^72^. In the present study, *pds5a syn4* double mutants lose TAD-like domains and revert contact decay to wild-type-like profiles (Fig. 2b). These findings indicate that, among somatic kleisins, SYN4-containing cohesin complexes are the principal engines for loop formation that becomes visible once PDS5A-mediated unloading is relaxed.

The domain-swap and Tudor-mutant analyses further refine how PDS5A differs functionally from other *Arabidopsis* PDS5 proteins. Chimeric proteins in which the central Armadillo array or the C-terminal Tudor-containing region of PDS5A are replaced by the corresponding segments of PDS5B fail to rescue the *pds5a* Hi-C phenotype, whereas a construct lacking the N-terminal Armadillo block largely retains TAD-like domain suppression (Fig. 3). In parallel, in the Tudor-mutant line, loss of H3K4me1 binding severely reduces PDS5A chromatin association yet fully restores wild-type-like contact decay and suppresses TAD-like domains, demonstrating that PDS5A-mediated unloading of cohesin and repression of loop extension do not require direct tethering to H3K4me1-marked chromatin (Fig. 4). These observations indicate that the PDS5A-specific central Armadillo architecture together with its C-terminal region, rather than its H3K4me1-binding capacity, are critical for limiting cohesin-driven chromatin extrusion, providing a mechanistic basis for the uniquely strong unloading activity of PDS5A among the plant PDS5 homologs.

Comparing the transcriptome of different *pds5a* tissues revealed that differentially expressed genes do not preferentially reside inside, outside, or at the borders of TAD-like domains, and insulation scores at regions harboring DEGs remain largely unchanged relative to the genome background (Fig. 1). This echoes animal studies where global TAD weakening or reshuffling yields only limited transcriptome changes, reinforcing that TADs are not universally required for baseline expression programs ^21,22,47^. However, targeted genetic perturbation of distal loop anchors reveals that specific promoter–promoter contacts created in *pds5a* can function as noncanonical cis-regulatory modules (Fig. 6). T-DNA insertions tens of kilobases away from upregulated genes abrogate *pds5a*-dependent transcriptional activation without disrupting local loop structures. These insertions alter gene expression only in the context of *pds5a*-dependent chromatin loops, implying that extrusion-driven juxtaposition enables distal promoters to act analogously to enhancers, extending functional communication well beyond the typical sub-kilobase enhancer–promoter distances characterized in *Arabidopsis*. These observations support a view in which the primary role of extrusion-mediated domains in plants does not to globally change transcription but to modulate the spectrum of specific long-range regulatory elements. In wild-type, strong PDS5A activity keeps extrusion short-ranged, constraining most cis-regulatory communication within local promoter neighborhoods, whereas in *pds5a*, extended loops expose genes to additional distal promoters that can either activate or repress transcription depending on their regulatory content.

In conclusion, our findings place PDS5A and SYN4-containing cohesin as central components of a plant-specific chromatin extrusion module that generates TAD-like structures and cis-regulatory loops when cohesin unloading is relaxed. Our work helps reconcile the apparent discrepancies between small- and large-genome plants and aligns plant 3D genome organization with conserved loop extrusion principles observed across eukaryotes ^8^. Future efforts to manipulate PDS5A and SYN4 in other plant species and developmental contexts should clarify the extent to which extrusion-driven loops are exploited for adaptive gene regulation, and whether tuning cohesin unloading can be used to engineer chromatin topology and transcriptional outputs in crops.

## Materials and Methods

### Plant materials and growth conditions

The wild-type reference used in this study is the Arabidopsis thaliana accession Columbia (Col-0). The T-DNA insertion mutant lines *pds5a-1* (*SALK_114556*), *wapl1-1*(*SALK_076791*), *wapl2-1*(*SALK_127445*), *syn2* (*SALK_044851*), *syn4* (*SALK_076116*), *SALK_020134*, *SALK_059642*, *SALK_041689* were obtained from the Nottingham Arabidopsis Stock Centre.

After surface-sterilization Arabidopsis seeds were sown on vertical half-strength Murashige and Skoog (1/2 MS) media plates (1% sucrose; 0.3% Phytagel). Following a 72 h stratification at 4°C the plants were grown in growth chambers (MLR-352-PE from PHCbi) at 21°C in long day condition (16 h light/8h dark). Heat stress treatment was performed by transferring 11-day-old plants to a growth chamber set at 37°C by day; the other parameters remaining identical. Heat-stressed plants were harvested three days later for subsequent analyses.

### Plasmid Construction

Genomic DNA containing the fragments of *PDS5A* and *PDS5B*, which were used to create the chimeric fusion proteins (Fig. 3a) were amplified with oligos listed in Supplementary Table 5. The respective fragments were cloned into the *pFK206* vector ^42^ via Gibson assembly, resulting in chimeric fusion proteins with a tandem HA-tag at the C-terminus. The construct for *PDS5A:2HA*^3A^ was based on *PDS5A:2HA*, which we reported previously ^42^. Mutagenesis PCR was performed using oligos listed in Supplementary Table 5 to introduce mutations at three amino acids in the Tudor domain of PDS5A (W1377A, Y1384A, Y1403A).

### In situ Hi-C

*in situ* Hi-C libraries were prepared essentially following a previously described protocol ^73^. Two independent Hi-C libraries were generated per sample, each starting from approximately 0.3 g of crosslinked material, homogenized for nuclei extraction. The isolated nuclei were resuspended in 150 µl 0.5% SDS and divided into three aliquots. After incubation at 62°C for 5 minutes, 145 µl water and 25 µl 10% Triton X-100 were added to quench SDS, followed by incubation at 37°C for 15 minutes. Chromatin digestion was carried out overnight at 37°C with 50 U DpnII (NEB) per tube. The next day, DpnII was inactivated at 62°C for 20 minutes. Subsequently, sticky ends were filled in using 1 µl each of 10 mM dTTP, dATP, dGTP, 10 µl 1 mM biotin-14-dCTP, 29 µl water, and 40 U Klenow fragment (Thermo Fisher) at 37°C for 2 hours. Proximity ligation was performed by adding 663 µl water, 120 µl 10× blunt-end ligation buffer (300 mM Tris-HCl, 100 mM MgCl_2_, 100 mM DTT, 1 mM ATP, pH 7.8), and 40 U T4 DNA ligase (Thermo Fisher) at room temperature for 3 hours. The resulting nuclei were collected by centrifugation, resuspended in 650 µl SDS buffer (50 mM Tris-HCl, 1% SDS, 10 mM EDTA, pH 8.0), and treated with 10 µg proteinase K (Thermo Fisher) at 55°C for 30 minutes, followed by reverse crosslinking in 30 µl 5 M NaCl at 65°C overnight.

DNA was isolated, treated with RNase A at 37°C for 30 minutes, and purified. Hi-C DNA (1 µg) was adjusted to 130 µl in TE buffer (10 mM Tris-HCl, 1 mM EDTA, pH 8.0) and sheared to fragments with an average size of 500 bp using a Q800R3 sonicator (QSONICA; 25% amplitude, 15 s on/off pulses, total 4.5 min). Fragments larger than 300 bp were recovered with Ampure beads. For biotin removal from unligated ends, DNA in a 50 µl reaction was incubated with 0.5 µl of 10 mM dTTP, 0.5 µl 10 mM dATP, and 5 U T4 DNA polymerase at 20°C for 30 minutes, followed by another Ampure cleanup. End repair and adaptor ligation were performed with the NEBNext® Ultra™ II DNA Library Prep Kit (NEB). Streptavidin C1 beads (Invitrogen) were used for affinity purification, and the libraries were PCR-amplified prior to sequencing with 2 × 150 bp reads on an Illumina Novaseq 6000 platform.

### Micro-C

Plant tissues were fixed and nuclei were isolated as described ^39^. Following the second crosslinking step with disuccinimidyl glutarate, nuclei were washed in MBuffer (10 mM Tris-HCl, pH 8.0, 50 mM NaCl, 5 mM MgCl₂, 1 mM CaCl₂, 0.1% Triton X-100, and 1× EDTA-free Complete protease inhibitor cocktail; Roche). MNase titration assays and Micro-C-XL library preparation were then performed essentially as described ^74^ with the following adaptations. First, all micrococcal nuclease (MNase) digestions were carried out in MBuffer instead of the original digestion buffer. MNase digestion profiles were evaluated on agarose gels to select conditions that yielded predominantly mono-and di-nucleosomal fragments. For library preparation, chromatin was digested under the selected conditions, end-repaired and ligated in situ, and crosslinks were reversed. Di-nucleosomal DNA was then size-selected using the PippinHT system (Sage Science) according to the manufacturer’s instructions. Finally, sequencing libraries were prepared from the size-selected DNA using the NEBNext Ultra II DNA Library Prep Kit (NEB).

### Capture Micro-C

BAC plasmids (F8M21, T21P5, and T17B22) served to generate probes for capture Micro-C, which were labeled with biotin-16-dUTP (Roche) via nick translation. Labeled probes were pooled into 20 µl hybridization mix (Twist Bioscience) for a final concentration of 0.5 ng/µl for each BAC. Separately, 300 ng of each indexed Micro-C library (NEBNext Ultra II DNA Library Prep Kit) was mixed with 0.6 µl NEXTFLEX Universal blockers in 12 µl library solution. Hybridization and streptavidin pull-down followed the Twist Target Enrichment Hybridization v2 Protocol: after denaturation at 95°C and cooling, probe and library solutions were combined and topped with Hybridization Enhancer. Hybridization was performed at 70°C for 16 h; products were captured with Dynabeads MyOne Streptavidin C1, washed at 68°C and 48°C, eluted in Tris-HCl (pH 8.0), and PCR-amplified for 14 cycles with P5 and P7 primers.

### Hi-C and Micro-C data processing

Read mapping to the TAIR10 reference was performed using Bowtie 2 (v2.2.4) ^75^, followed by PCR duplicate removal and filtering. Hi-C reads were filtered according to our previously described procedures, which took into account the distribution of the restriction enzyme recognition site (5’-GATC-3’) to ensure that only *bona fide* Hi-C ligation products were retained. In contrast, all inter-chromosomal Micro-C reads were retained, as were all intra-chromosomal reads for which the mapped distance between read1 and read2 exceeded 500 bp in the reference genome. Hi-C and Micro-C reads filtering statistics for all samples are provided in Supplementary Table 1. Iterative matrix correction for map normalization was performed with the HiTC R package ^76^; matrix normalization was stopped when the *eps* value fell below 1 × 10⁻⁴. Additionally, hic files for interactive visualization were generated using Juicer, on the filtered, informative Hi-C and Micro-C reads ^77^.

### Chromatin insulation, TAD-like domains and chromatin loop annotation

Insulation scores were calculated according to a previous study ^42^. We used Hi-C maps normalized with 5 kb bins, and intrachromosomal contacts up to 100 kb to compute insulation scores. Potential “insulated regions” were identified by searching for regions with local insulation score minima, which was done by using the peak function from the splus2R package in R with span=10. In this study, we annotated the regions with insulation scores lower than -0.25 and overlapping with insulation score local minima as “insulated regions”. A list containing insulation scores and insulated region annotation is available in Supplemental Table 2.

TAD-like domains were annotated using the Hi-C contact map from *pds5a* leaf tissues, normalized at 5 kb resolution. The arrowhead algorithm, which we previously applied to similar chromatin features in various plant species, was used for domain identification ^29,42,73^. For this study, the TAD score matrix cutoff was set to 0.95, candidate TADs were required to contain at least 6 filtered pixels, and the minimum TAD score at domain borders was set to 1.10.

Chromatin loops were called using SIP (v1.6.5.) ^78^ with the following parameters: *“-norm KR -g 1.5 -min 2.0 -max 2.0 -mat 5000 -d 6 -res 1000 -sat 0.01 -t 2800 -nbZero 6 -factor 1 -fdr 0.01 -del true -cpu 1 -isDroso false*” on Micro-C maps of *pds5a* normalized at 1 kb resolution.

For both types of annotation, the following pericentromeric regions were excluded: Chr1 (11,500,001-17,700,000), Chr2 (1,100,001-7,200,000), Chr3 (10,300,001-17,300,000), Chr4 (1,500,001-6,300,000), and Chr5 (9,000,001-16,000,000).

### Stripe annotation

Stripes were detected using the Zebra algorithm ^18^. For each chromosome, both ICE-normalized and raw contact matrices were used. Contacts between loci separated by more than 400 kb were excluded. For ICE-normalised matrices, all zero values were also set to NA. For each pixel (*i,j*), in each matrix of size *n × n*, the following metrics were computed:

1. Sum of raw counts and median of ICE-normalized values in a local window: To quantify local interaction strength around the given pixel (*i,j*), the sum of raw contact counts (Sum_Local_) within a window of fixed size was calculated:

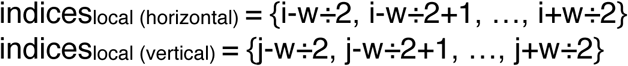

where *w* is the window size and ÷ denotes integer division. The sum of values in the row or column over these indices represents the horizontal or vertical local interaction signal, respectively. If the pixel (*i,j*) had less than 5 reads, the local sum was set to NA. Additionally, for the ICE-normalised matrix, the median value across the same local window is computed as Median_local_. For stripe detection in 1 kb matrices, the window size w was set to 5 pixels.
2. Median of ICE-normalized values of neighbourhood: To estimate a background signal away from the pixel (*i,j*), the median ICE-normalised values was calculated in four neighbourhood regions: top, bottom, left, and right (Median_direction_). Each region was defined using a fixed window *f* and a gap *g* between the pixel (*i,j*) and the neighbourhood defined as:

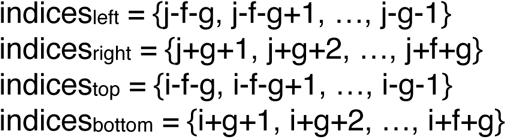

where *j* and *i* denoted column and row indices, respectively. For stripe detection in 1 kb matrices, the window size f was set to 4 pixels and the gap g to 2 pixels.
3. Bias and expected value: Next, the horizontal and vertical biases for both horizontal and vertical local regions were defined as:

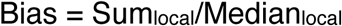 Which were used to compute expected values for individual direction (top, bottom, left, and right) as:

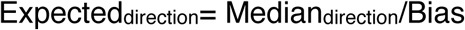

with top and bottom using the horizontal bias, and left and right using the vertical bias.
4. Statistical testing: Poisson tests were performed for each pixel to evaluate whether the observed local contact count is significantly greater than the expected value derived from its neighbourhood. The tests were applied as follows: For testing for downstream stripes, the observed value was the horizontal *Sum_local_*, and two tests were performed against *Expected_top_* and *Expected_bottom_*. For upstream stripes, the observed value was the vertical *Sum_local_*, and two tests were performed against *Expected_left_* and *Expected_right_*. A pixel was considered potentially part of a stripe if both tests in the corresponding directions satisfied the following thresholds: FC (Fold change) > 1.1 and p-value < 0.005. Pixels meeting these conditions were assigned a value of 1 in a binary matrix of size *n* × *n*. All other pixels were assigned 0.

Next, stripe regions were identified from the binary enrichment matrix by scanning along each row (for downstream stripes) and each column (for upstream stripes) to find consecutive runs of pixels with value 1. Run-length encoding (RLE) was applied to each row or column to identify these runs. Runs were selected as candidate stripes if their length met or exceeded 5 pixels, and their row (or column) index and start/end positions were recorded. Adjacent candidate stripes on the same row or column were merged if separated by a gap less than or equal to 3 pixels, with merging performed by extending the end position of the first stripe. After merging, only stripes with a total length above 11 pixels were retained. For vertical stripes, those located in adjacent columns, and for horizontal stripes, those in adjacent rows, were grouped if their positions differed by two bins or fewer. Each group was then consolidated by recording the minimum and maximum values for row/column, start, and end positions. Finally, stripes anchored within the following pericentromeric regions were excluded: Chr1 (11,500,001-17,700,000), Chr2 (1,100,001-7,200,000), Chr3 (10,300,001-17,300,000), Chr4 (1,500,001-6,300,000), and Chr5 (9,000,001-16,000,000).

### De novo motif analysis

DNA motifs within pds5a chromatin loop were identified by first extracting sequences from accessible chromatin regions located (ATAC-seq peaks) within the loop anchors. For this, the two peak sets from *pds5a* replicate 1 and 2 were merged based on overlapping peaks with the “findOverlaps” function from the R package GenomicRanges ^79^. Peaks within stripe anchors were extracted and only peaks fully contained within an anchor region were retained for downstream analysis. Sequences corresponding to these filtered peaks were then extracted from the TAIR10 reference genome FASTA file.

The sequences were submitted to MEME for de novo motif discovery ^80^ with the following settings: *“-dna -oc. -nostatus -time 14400 -mod anr -nmotifs 3 -minw 4 - maxw 12 -objfun classic -revcomp -markov_order 0*”.

### Chromatin immunoprecipitation and data analyses

Arabidopsis shoot tissue was fixed under vacuum for 25 minutes at room temperature in 2 mM DSG in PBS buffer supplemented with 0.01% Triton X-100. Subsequently, formaldehyde stock solution was added to achieve 1% final concentration, and the samples were further incubated in vaccum for 25 mins. To quench the fixation, the solution was replaced with 0.15 M glycine in PBS buffer, and samples were incubated under vacuum for an additional 10 minutes at room temperature. Approximately 0.5 g of the fixed tissue was then homogenized and resuspended in nuclei isolation buffer (20 mM HEPES, pH 8.0; 250 mM sucrose; 1 mM MgCl_2_; 5 mM KCl; 40% glycerol; 0.25% Triton X-100; 0.1 mM PMSF; 0.1% 2-mercaptoethanol). The isolated nuclei were pelleted and subsequently resuspended in 0.5 mL sonication buffer (10 mM potassium phosphate, pH 7.0; 0.1 mM NaCl; 0.5% sarkosyl; 10 mM EDTA). Chromatin was fragmented by sonication using a QSONICA Q800R3 sonicator, resulting in an average fragment size of approximately 500 bp. Fifty microlitres of 10% Triton X-100 were added to the sonicated chromatin, and the mixture was further mixed with an equal volume of IP buffer (50 mM HEPES, pH 7.5; 150 mM NaCl; 5 mM MgCl_2_; 10 µM ZnSO_4_; 1% Triton X-100; 0.05% SDS) and incubated with Pierce™ Anti-HA magnetic beads (Thermo Fisher) for 2 hours at 4°C. Following immunoprecipitation, the beads were washed at 4°C as follows: twice with IP buffer, once with IP buffer containing 500 mM NaCl, and once with LiCl wash buffer (0.25 M LiCl; 1% NP-40; 1% sodium deoxycholate; 1 mM EDTA; 10 mM Tris-HCl, pH 8.0), each for 3 minutes. Beads were then briefly washed with TE buffer (10 mM Tris-HCl, pH 8.0; 1 mM EDTA). For elution, the beads were resuspended in 200 µL elution buffer (50 mM Tris-HCl, pH 8.0; 200 mM NaCl; 1% SDS; 10 mM EDTA) and incubated at 65°C for 6 hours, followed by treatment with Proteinase K at 45°C for 1 hour. DNA was subsequently purified using the MinElute PCR Purification Kit (QIAGEN) and was either analyzed by qPCR or used to generate sequencing libraries with the NEBNext® Ultra™ II DNA Library Prep Kit (NEB). The libraries were sequenced with 2 × 150 bp reads on an Illumina Novaseq 6000 platform.

ChIP-seq reads were mapped to the TAIR10 genome using Bowtie 2 ^75^. H3K4me1 ChIP-seq analysis was conducted via processing the raw sequencing data described in a previous study ^61^, and peak calling was done with MACS2 v.2.1.1 ^81^ using parameters *"-g 1.2e8 --bw 450 --keep-dup=1*".

### Gene expression analyses

RNA sequencing (RNA-Seq) was performed with two replicates per sample. Total RNA was extracted from unfixed seedlings, roots, and germinating seeds by mechanically disrupting the plant material in liquid nitrogen, followed by RNA extraction using the RNeasy Plant Mini Kit (Qiagen), according to the manufacturer’s instructions. For each sample, 500 ng of total RNA was treated to remove residual genomic DNA by adding 0.5 μl 10x reaction buffer (100 mM Tris, pH 7.5, 25 mM MgCl_2_, 1 mM CaCl_2_), 0.5 μl DNase I, and 0.5 μl RNaseOUT (Thermo Scientific) and incubating at 37 °C for 30 min. The reaction was stopped by adding 0.5 μl 50 mM EDTA and incubating at 65 °C for 10 min. RNA-seq libraries were generated from the DNase-treated RNA using the NEBNext® Ultra™ RNA Library Prep Kit for Illumina (NEB E7770), NEBNext® Poly (A) mRNA Magnetic Isolation (NEB E7490) and NEBNext Multiplex Oligos for Illumina, following the manufacturer’s protocol at 0.5x reaction volume. The libraries were sequenced with 2 × 150 bp reads on an Illumina Novaseq 6000 platform.

Sequencing reads were mapped to the TAIR10 reference genome using Tophat2 (v2.1.1) ^82^ with the “--no-novel-juncs” options. A count table with the number of reads per gene was generated by overlapping the aligned reads with TAIR10 gene annotations using the R package GenomicAlignments ^79^. Differential gene expression analysis was performed using the R package DESeq2 ^83^. Differentially expressed genes were identified based on a false discovery rate smaller than 0.05 and a fold change of greater than 2. Differentially expressed genes are listed in Supplementary Table 3.

For analysis of individual gene expression, total RNA was extracted from the samples using the RNeasy Plant Mini Kit (Qiagen) according to the manufacturer’s protocol. The RNA was subsequently treated with DNase I (Thermo Scientific) to eliminate residual genomic DNA. Afterwards, first-strand cDNA was synthesized from the RNA using the qScriber™ cDNA Synthesis Kit (highQu). Quantitative RT-PCR was performed on an Applied Biosystems™ QuantStudio™ qPCR System (Thermo Scientific) employing qPCRBIO SyGreen Mix ROX (Lo-ROX) (PCR Biosystems) and gene-specific primers listed in Supplementary Table 5.

### Chromatin accessibility analyses

For ATAC-seq, approximately 0.1 g of fixed plant material was chopped on ice in 1 ml general purpose buffer (GPB; 0.5 mM spermine, 30 mM sodium citrate, 20 mM MOPS, 80 mM KCl, 20 mM NaCl, 0.5% (v/v) Triton X-100, pH 7.0) and filtered first through single-layer Miracloth (Merck Millipore, with pore size 22-25 μm) and then through a cell strainer. DAPI was added to a final concentration of 600 nM, and 50,000 diploid nuclei were sorted into NIB by fluorescence-activated cell sorting (FACS) based on DAPI signal, using a Bio-Rad S3e cell sorter.

ATAC-seq was performed with two biological replicates. Nuclei were first incubated at 60 °C for 5 min and centrifuged at 1000 x g at 4 °C for 5 min. To the nuclei pellet, 10 μl TD, 1 μl TDE1 (Nextera® DNA Library Prep kit) and 9 μl water were added and the reaction was incubated at 37 °C for 30 min. The reaction was then adjusted to 200 μl with SDS buffer (50 mM Tris, pH 8.0, 1% (w/v) SDS, 10 mM EDTA) and 8 μl 5 M NaCl was added. After incubating overnight at 65 °C, DNA was purified using the Qiagen Mini-purification kit (Qiagen). To prepare DNA for sequencing, 2 μl of DNA were used as template for PCR using the NEBNext® High-Fidelity 2X PCR Master Mix (NEB) and Nextera index primers (Illumina). The following PCR protocol was applied: 72 °C for 5 min, followed by initial denaturation at 98 °C for 20 s, 12 cycles of denaturation at 98 °C for 10 s, annealing at 63 °C for 30 s, and extension at 72 °C for 1 min, finally 72 °C for 1 min. DNA was then size-selected using magnetic beads (SpeedBead Magnetic Carboxylate Modified Particles, Cytiva) to recover fragments between 200 and 500 bp.

ATAC-seq reads mapping was performed according to our previous study ^84^. BigWig and count files were generated using deepTools (v3.5.5) with “bamCoverage” and BEDTools (v2.31.1) with the “multicov” function, respectively. Accessible peaks were defined from ATAC-seq data using MACS2 (v2.1.1-python-2.7.12) peak calling with a genome size of 1.2 × 10^8^ and duplicate reads limited to one (--keep-dup=1).

### Protein extraction and western blot

Total protein was extracted from 14-day-old aerial tissue using TRIzol-chloroform and subsequently dissolved in a buffer containing 88 mM Tris-HCl (pH 6.8), 4.4 M urea, 2.2% SDS, 10 mM DTT, and 11% glycerol. The protein samples were analyzed by western blot and HA-tagged proteins were detected by anti-HA-HRP (1:2500; sc-7392, Santa Cruz Biotechnology). The chemiluminescent signals were detected with HRP substrate (Takara) in the iBright CL750 Imaging System.

### Isolation of PDS5A-interacting proteins

Shoot tissue of 14-day-old plants was fixed for 30 min under vacuum with 1% formaldehyde in MC buffer (10 mM potassium phosphate, pH 7.0, 50 mM NaCl, 0.1 M sucrose). The fixative was replaced with 0.15 M glycine in MC buffer, wherein the plant material was incubated under vacuum for an additional 10 minutes. Approximately 0.5 g fixed tissue was homogenized in nuclei isolation buffer (20 mM Hepes, pH 8.0, 250 mM sucrose, 1 mM MgCl2, 5 mM KCl, 40% glycerol, 0.25% Triton X-100, 0.1 mM PMSF, 0.1% 2-mercaptoethanol). The homogenate was filtered through double-layered miracloth (Milipore). The isolated nuclei were washed in NIB and subsequently resuspended in 0.5 ml sonication buffer (10 mM potassium phosphate, pH 7.0, 0.1 mM NaCl, 0.5% sarkosyl, 10 mM EDTA). The samples were sonicated five times for 10 seconds, with a resting period of 50 seconds between each sonication with the sonicator (QSONICA Q800R3) set at 40% of full capacity. After the addition of 50 µL 10% Triton X-100 the sonicated samples were mixed with an equal volume of IP IP buffer (50 mM HEPES, pH 7.5; 150 mM NaCl; 5 mM MgCl_2_; 10 µM ZnSO_4_; 1% Triton X-100; 0.05% SDS) and Pierce™ Anti-HA magnetic beads (Thermo Fisher). The IP was carried out at 4°C for 30 minutes and the beads subsequently collected and washed as described in chromatin immunoprecipitation. Finally, the beads were resuspended in TE buffer, mixed with 5x Laemmli Loading Buffer, and boiled at 95°C for 10 minutes. The proteins were separated on a 10 % SDS-PAGE gel and submitted to the Core Facility Hohenheim for analysis by mass spectrometry.

## Supporting information

HiC and MicroC libraries

TADs, Stripes, and chromatin loops in pds5a

DEG list

proteomics data

oligo sequences

## Data availability

Short read data of in situ Hi-C, micro-C, ChIP-seq, ATAC-seq, and RNA-seq are publicly available at NCBI Sequence Read Archive under accession number PRJNA1392550.

Large datasets, such as normalized Hi-C and micro-C matrices, BigWig ChIP-seq and ATAC-seq track files, are available in the figshare repository. They are accessible with the following link with Digital Object Identifier (DOI) 10.6084/m9.figshare.30814679

## Code availability

All scripts used in this study are available upon request.

## Funding

This work was supported by the Deutsche Forschungsgemeinschaft (LI 2862/8 and LI 2862/11) and intramural funding from the University of Hohenheim.

## Acknowledgements

We thank computing support by the High Performance and Cloud Computing Group at the Zentrum für Datenverarbeitung of the University of Tübingen, the state of Baden-Württemberg through bwHPC and the German Research Foundation (DFG) through grant no. INST 37/935-1 FUGG. We acknowledge Jens Pfannstiel and Berit Würtz from the Core Facility Hohenheim for the support on mass spectrometry analysis. We thank Louisa Ford for assistance with gene expression measurement.

## Notes

### Competing Interest Statement

The authors have declared no competing interest.

