## supplemental Figs for "A molecular framework of chromatin extrusion in plants"

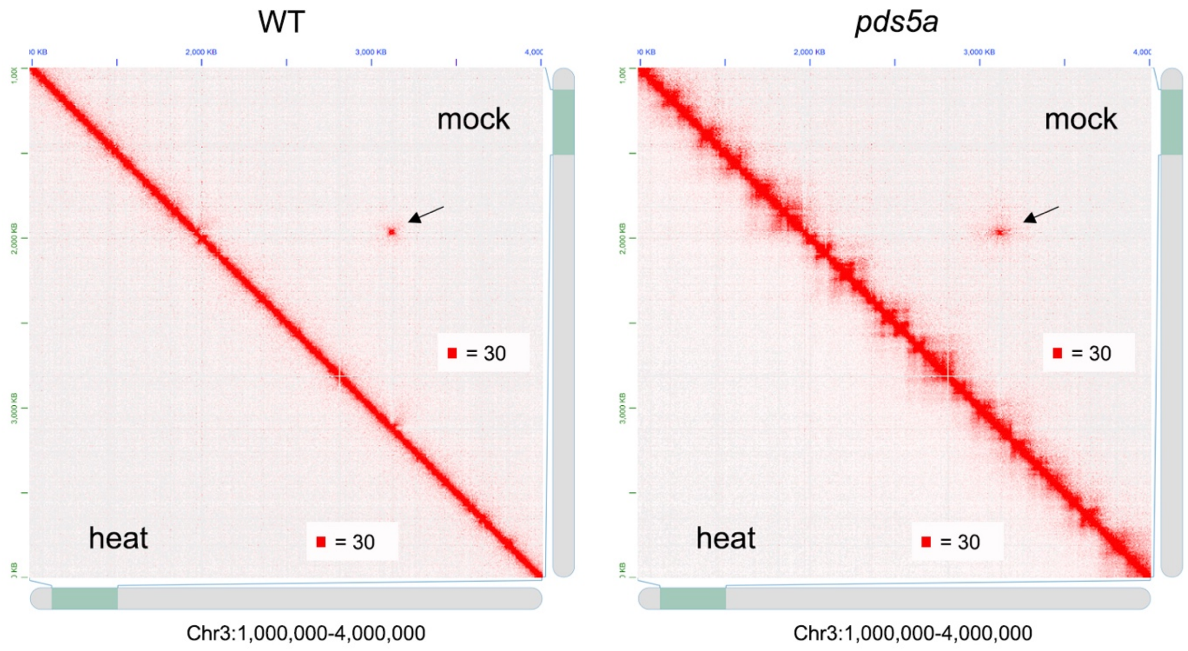


**Supplemental Figure 1 I Chromatin organization features exhibit distinct responses under heat stress.** The Hi-C contact maps, normalized to a 20 kb bin size, illustrate a 3 Mb segment of chromosome 3. Arrows highlight long-range chromatin interactions occurring between two KNOT ENGAGED ELEMENT regions, previously characterized in an earlier study[1].

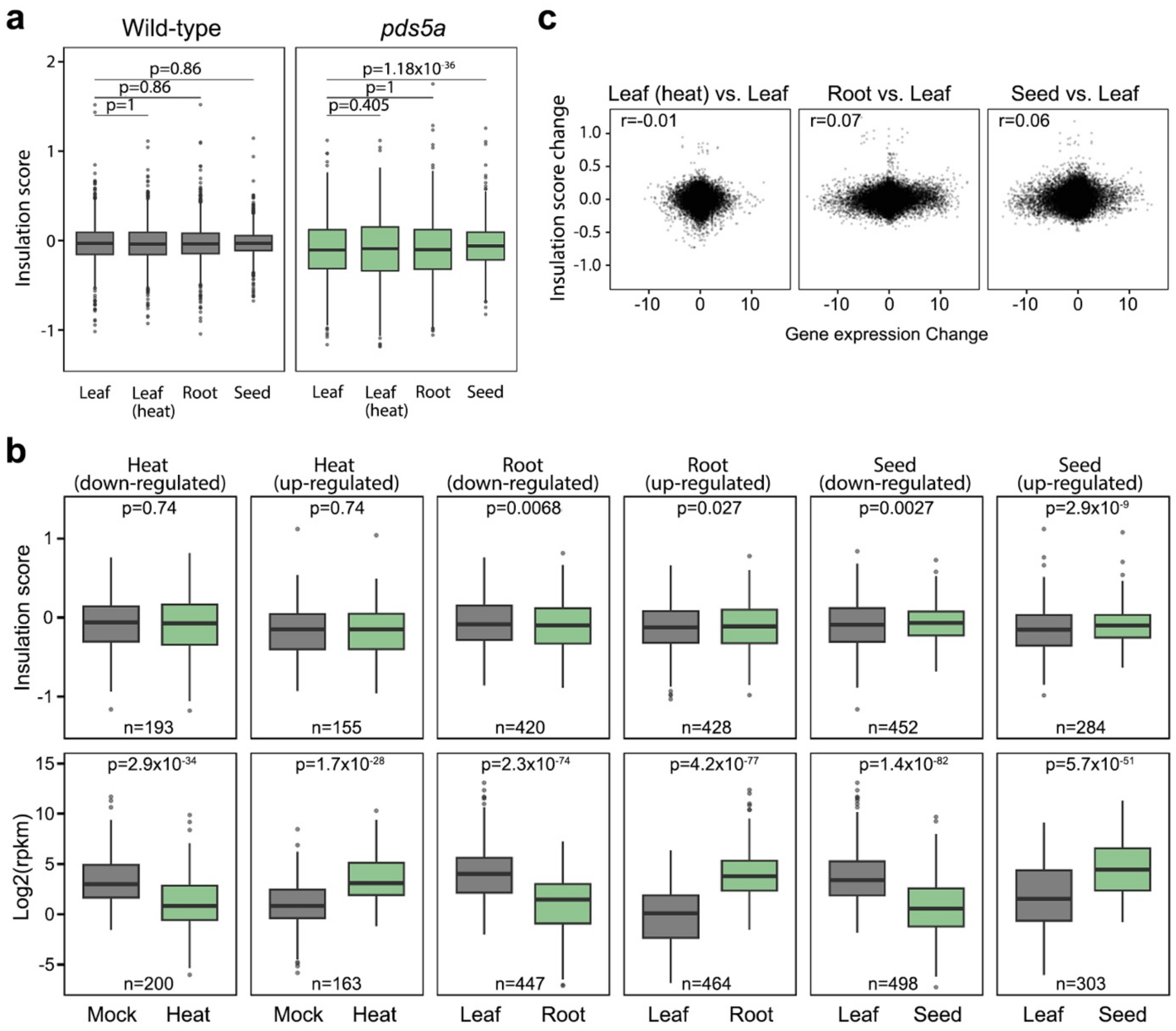


**Supplemental Figure 2 | Comparison of insulation scores at TAD boundaries in different *pds5a* tissues and conditions. a,** Boxplots show the distribution of insulation scores at TAD boundaries across tissues or conditions in wild-type (grey) and *pds5a* (green), with the box spanning the interquartile range (IQR; Q1-Q3), the horizontal line indicating the median, whiskers extending to 1.5×IQR, and outliers shown as individual points. P-values were calculated using paired Wilcoxon signed-rank tests and corrected using the Holm method within each genotype **b,** Boxplots of insulation scores (top panels) and log2(RPKM) expression values (bottom panels) at TAD boundaries overlapping transcription start sites (TSS) of differentially expressed genes (DEG) (|log2 fold-change| > 2 and p-value <= 0.05) in *pds5a*. Comparisons are shown for *pds5a* heat, root, and germinating seed samples versus mock. TAD boundaries were selected based on TSS overlap within a 2 kb bin. Sample sizes (n) indicate the number of selected TAD boundaries or genes per comparison. Statistical testing used paired, two-sided Wilcoxon signed-rank tests with Holm correction for insulation scores and gene expression values separately. **c,** Genome-wide correlation between change in insulation score and change in gene expression at all TSS. Scatterplots show Pearson's correlation coefficient (r) between log2 fold-change of gene expression and the corresponding change in insulation score at each TSS.

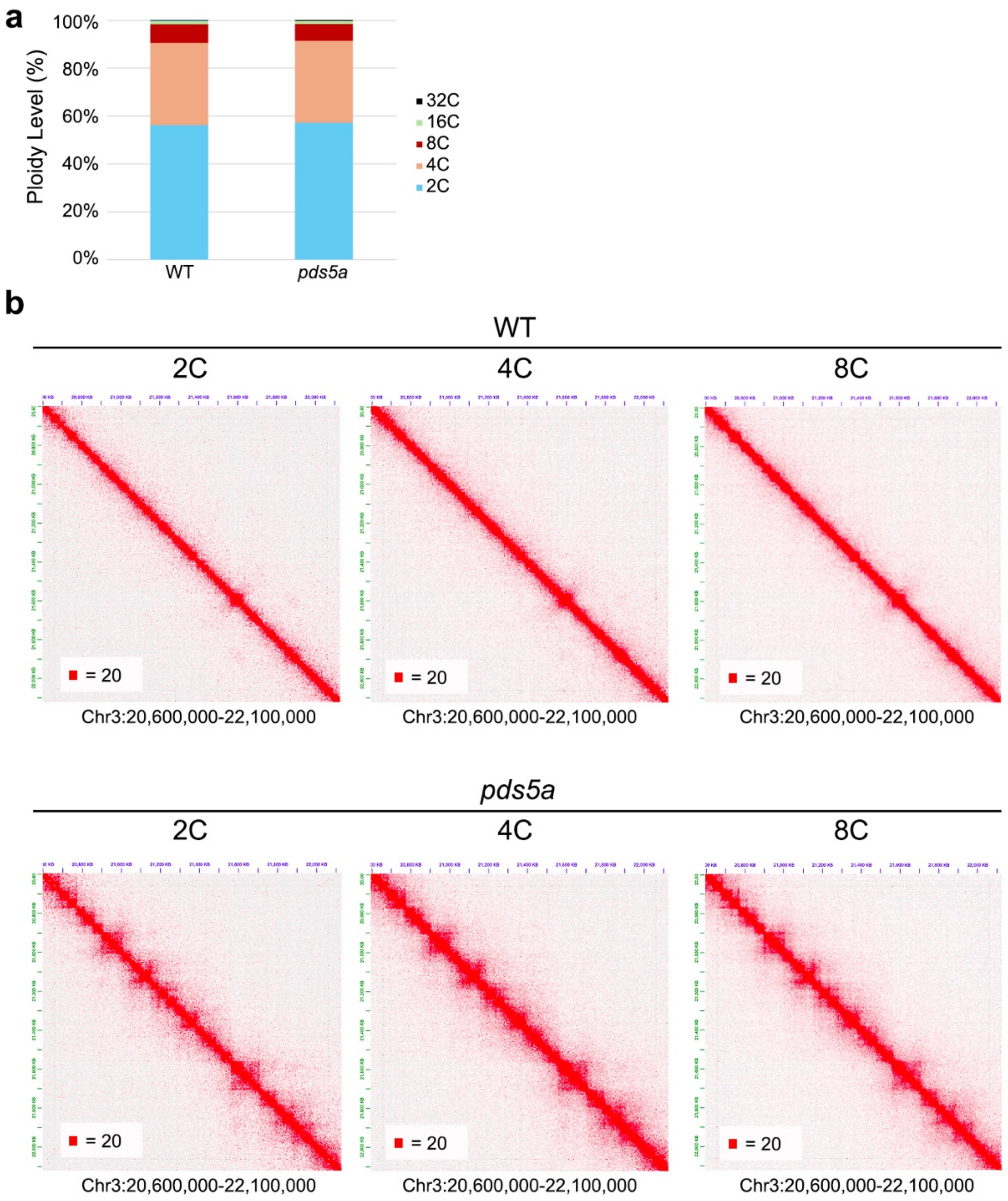


**Supplemental Figure 3 | TAD-like domains in pds5a are independent of endoreduplication cycles. a,** Endopolyploidy measurements in wild-type (WT) and *pds5a* seedlings. Nuclei were isolated from 14-day-old seedlings and the ploidy distribution was analyzed with the S3e Cell Sorter (Bio-Rad). **b,** TAD-like structures in *pds5a* are present in both nuclei with and without completing endoreduplication cycles. The Hi-C maps (normalized at a 5 kb bin size) show a 0.5 Mb genomic region from chromosome 3.

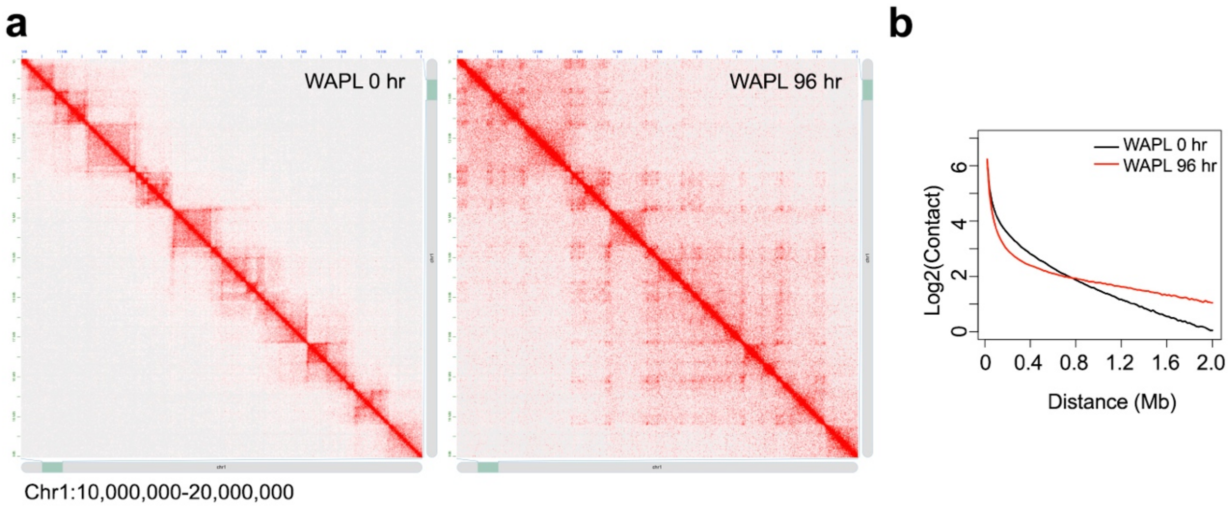


**Supplemental Figure 4 | Effects of WAPL depletion on chromatin organization in mammalian cells. a,** Comparison of Hi-C maps of a 10 Mb representative genomic region from mouse embryonic stem cells before (WAPL 0 h) and 96 hours after WAPL knockdown (WAPL 96 h). The Hi-C maps were reconstructed from a public dataset [2] and normalized at bin size of 20 kb. **b,** Genome-wide intrachromosomal-contact frequency plotted as a function of genomic distance.

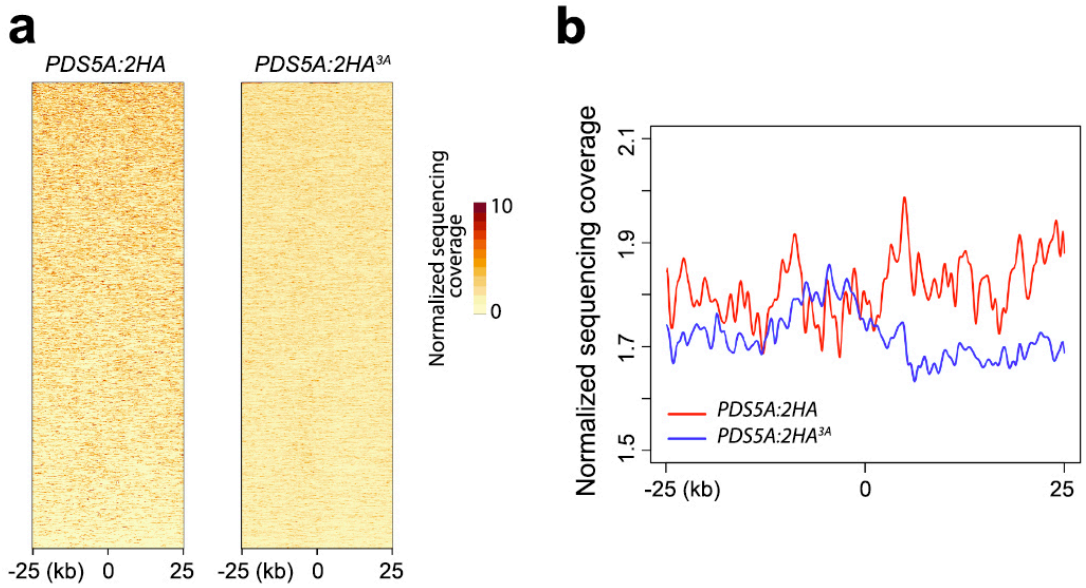


**Supplemental Figure 5 | Interactions between PDS5A and chromatin at TAD-like domain borders. a,b** Normalized ChIP-seq reads heatmap of PDS5A:2HA and the mutagenized PDS5A:2HA$^{3A}$ at individual (**a**) and averaged (**b**) TAD-like borders (n= 1208).

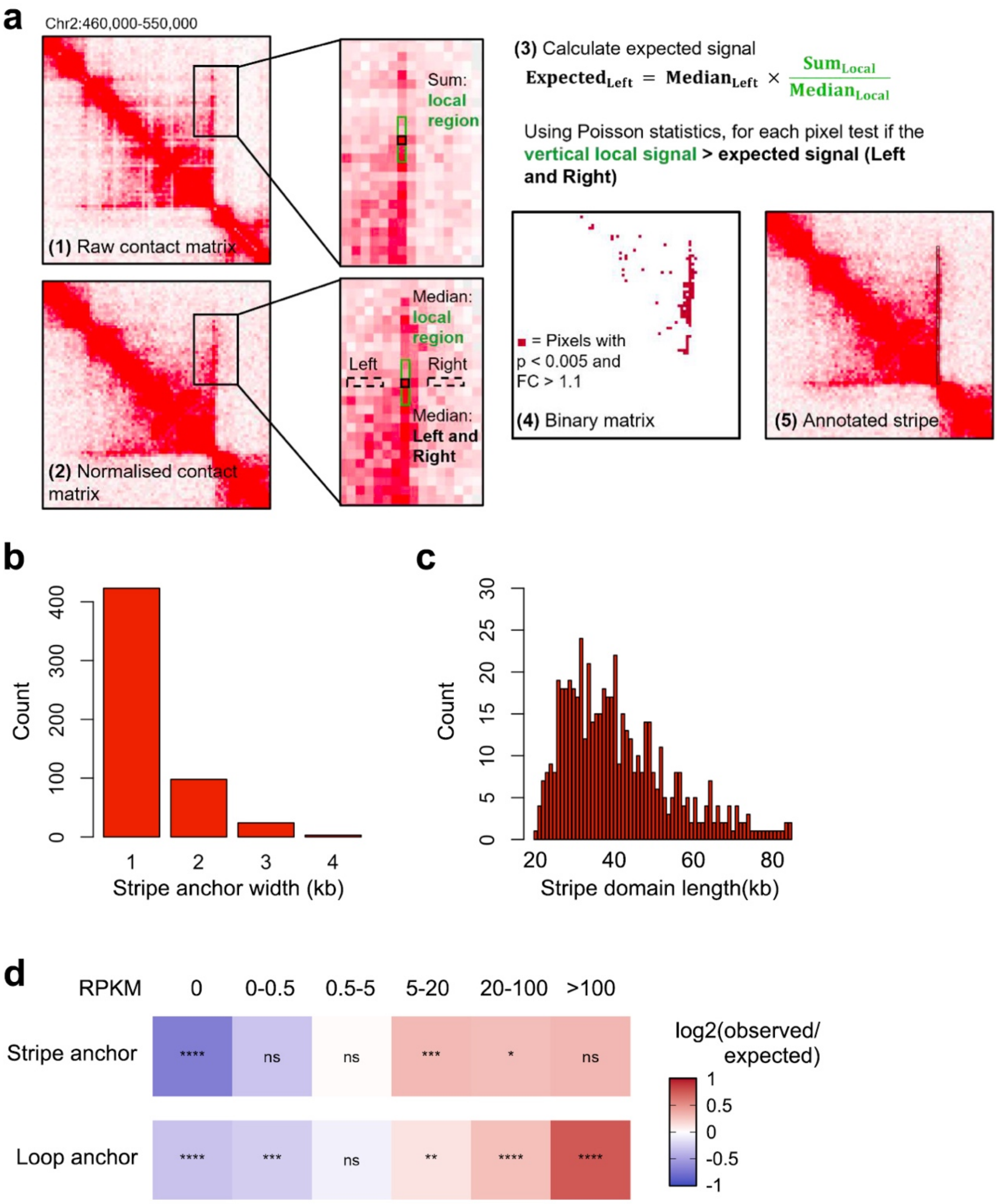


**Supplemental Figure 6 | Detection of architectural stripes in *pds5a* with the Zebra algorithm. a,** Computational approach for stripe detection, illustrated for an upstream stripe. (1) For each pixel located at a maximum distance from the diagonal, a vertical local region is defined, including the focal pixel and its two adjacent pixels (green), and the raw contact values are summed. (2) Additionally, two flanking neighbourhood regions (Left and Right; dashed boxes), located two pixels away from the local region, are extracted. Median contact values are computed for the local and neighbourhood regions using ICE-normalised Hi-C data. (3) The expected background signal is calculated, and Poisson statistics are applied to test whether the local signal read count exceeds the expected background value. (4) Each pixel yields two fold-change (FC) values and two associated p-values (Left and Right). Pixels satisfying both FC > 1.1 and p < 0.005 in both comparisons are retained and clustered to obtain stripes. (5) Stripes are annotated as linear regions extending from the anchor point to the most distal retained pixel. For downstream stripes, the same analysis is performed using a horizontal local region and top and bottom neighbourhood zones. **b,** Distribution of stripe anchor widths. **c,** Distribution of stripe domain lengths measured from the most distant contact from the anchor. **d,** Enrichment of gene expression categories in stripe anchors and chromatin loop anchors relative to genomic background. Genes outside pericentromeric regions were binned by average RPKM values from pds5a leaf samples into six categories: non-expressed (RPKM = 0; n = 5,403), very low (0-0.5; n = 3,182), low (0.5-5; n = 6,718), moderate (5-20; n = 6,504), high (20-100; n = 3,394), and very high (>100; n = 875). Colours indicate log2(observed/expected) enrichment, with asterisks denoting statistical significance from bin-wise chi-squared goodness-of-fit tests with Benjamini-Hochberg correction (**** p < 0.0001, ** p < 0.01, * p < 0.05, ns not significant).

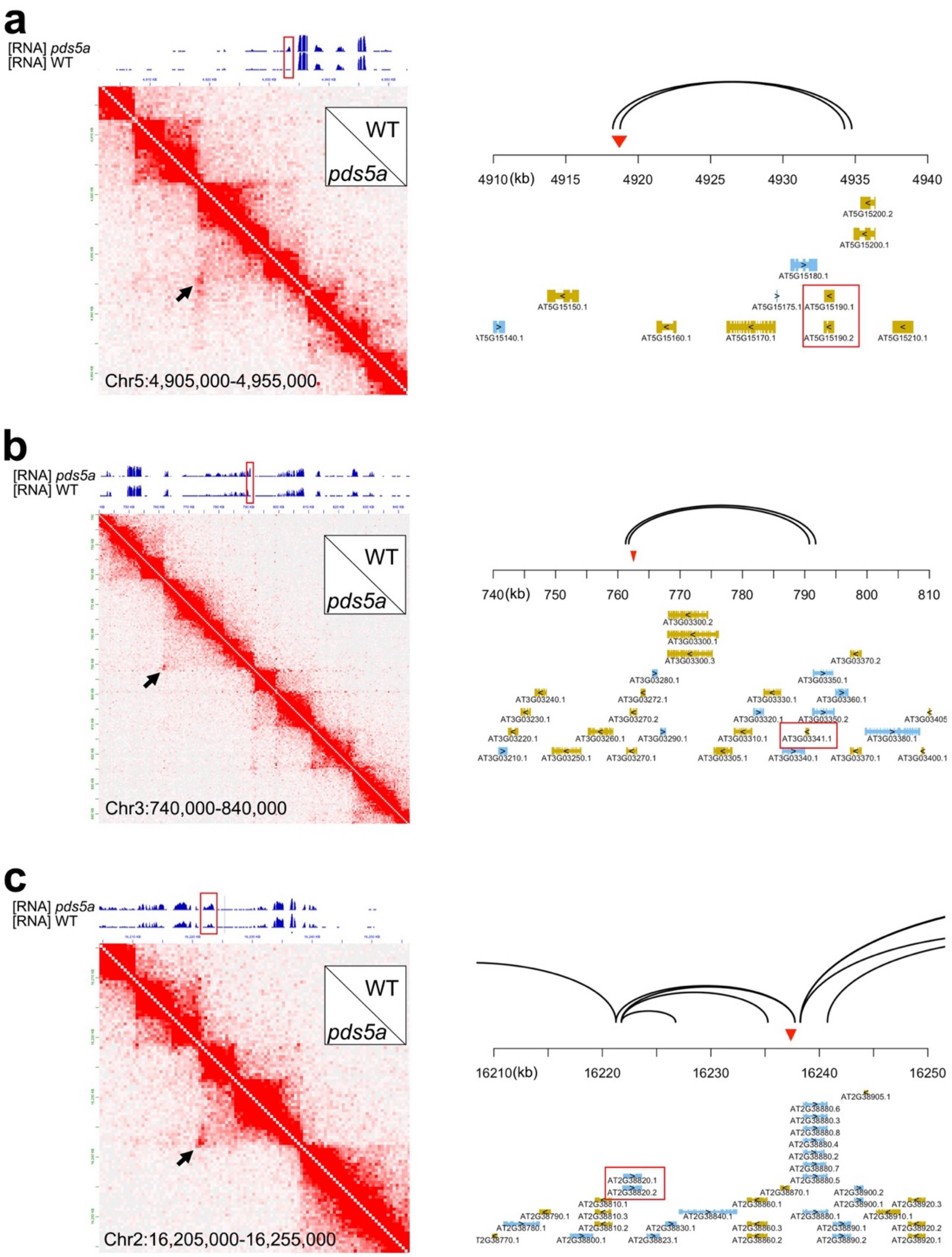


**Supplemental Figure 7 | Association between gene expression change and long-range chromatin contacts in pds5a. a-c,** Micro-C maps normalized to a 500 bp window size reveal the presence of prominent chromatin loops formed in *pds5a* (Arrows). RNA-seq coverage tracks above each map display gene expression in the corresponding genomic regions, with boxes indicating differentially expressed genes. For each panel, a magnified gene annotation is shown on the right: chromatin loops are represented by black arcs, and differentially expressed genes are highlighted by red rectangular frames. The red triangle marks the insertion site of the SALK line used in this study.

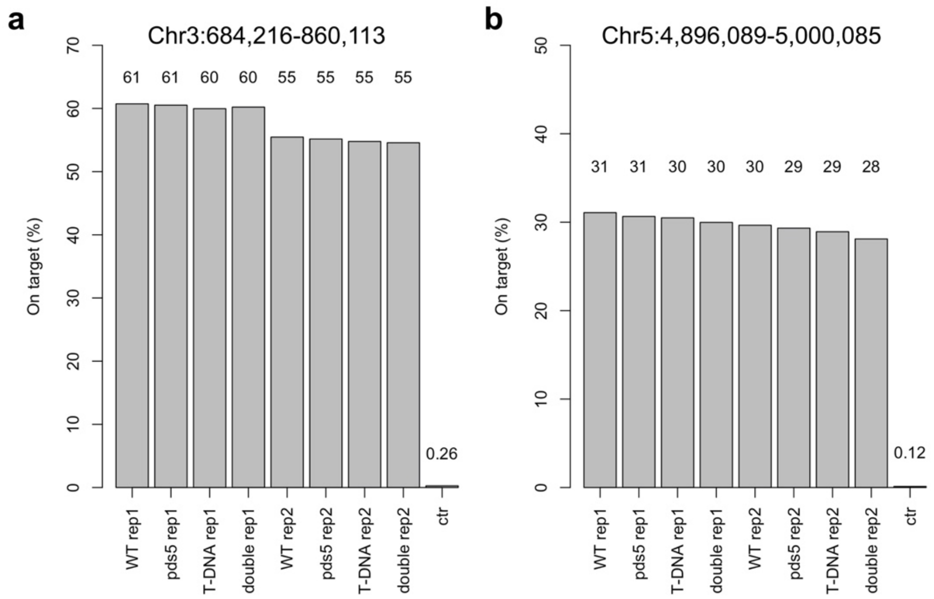


**Supplemental Figure 8 | Enrichment of Capture Micro-C libraries. a,b** Percentages of on-target Micro-C contacts in individual libraries, defined as reads overlapping with genomic regions covered by probes. ctr, a *pds5a* Micro-C library without hybridization-mediated capture, representing the baseline of Micro-C contacts concerning the region of interest.
